# Ubiquinone deficiency drives reverse electron transport to disrupt hepatic metabolic homeostasis in obesity

**DOI:** 10.1101/2023.02.21.528863

**Authors:** Renata L.S. Goncalves, Zeqiu Branden Wang, Karen E. Inouye, Grace Yankun Lee, Xiaorong Fu, Jani Saksi, Clement Rosique, Gunes Parlakgul, Ana Paula Arruda, Sheng Tony Hui, Mar Coll Loperena, Shawn C. Burgess, Isabel Graupera, Gökhan S. Hotamisligil

## Abstract

Mitochondrial reactive oxygen species (mROS) are central to physiology. While excess mROS production has been associated with several disease states, its precise sources, regulation, and mechanism of generation *in vivo* remain unknown, limiting translational efforts. Here we show that in obesity, hepatic ubiquinone (Q) synthesis is impaired, which raises the QH_2_/Q ratio, driving excessive mROS production via reverse electron transport (RET) from site I_Q_ in complex I. Using multiple complementary genetic and pharmacological models *in vivo* we demonstrated that RET is critical for metabolic health. In patients with steatosis, the hepatic Q biosynthetic program is also suppressed, and the QH_2_/Q ratio positively correlates with disease severity. Our data identify a highly selective mechanism for pathological mROS production in obesity, which can be targeted to protect metabolic homeostasis.

## Main Text

Mitochondria are the central organelles for processing nutrient-derived substrates, generating key metabolites vital for cell function. Intrinsic to this process is the generation of superoxide and hydrogen peroxide (referred to herein as mitochondrial reactive oxygen species or mROS), which are essential for cell signaling, but detrimental when generated in excess (*1*, *2*). The chronic metabolic stress imposed by overnutrition in obesity (*3*) has been linked to increased production of mROS, which is related to the development of metabolic pathologies including insulin resistance and type 2 diabetes (*4*–*8*). Yet, over the past two decades it has become clear that non-specific removal of these oxidants through systemic administration of broad and non-selective antioxidants is not effective in treating metabolic pathologies, as seen by the failure of large-scale clinical trials (*9*–*11*). Therefore, a more mechanistic approach directed to the specific source of mROS may be a safer and more effective intervention strategy (*12*). However, it has been challenging to determine the molecular mechanisms underlying excess mROS generation and its links to metabolic pathologies *in vivo* (*13*).

In fact, mROS production is not a single process. In intact cells and *in vivo*, the rate of mROS generation is the sum of the activity of at least 11 distinct sites associated with the electron transport chain (ETC) and matrix substrate oxidation (**Fig. 1C**). These 11 sites present substrate-specificity and show distinct capacities to generate superoxide (*14*). Notably, complex I is important not only because it is one of the major producers of superoxide, but also because it can operate bidirectionally, which has been associated with different physiological or pathological outcomes (*15*–*20*).

**Fig. 1.**
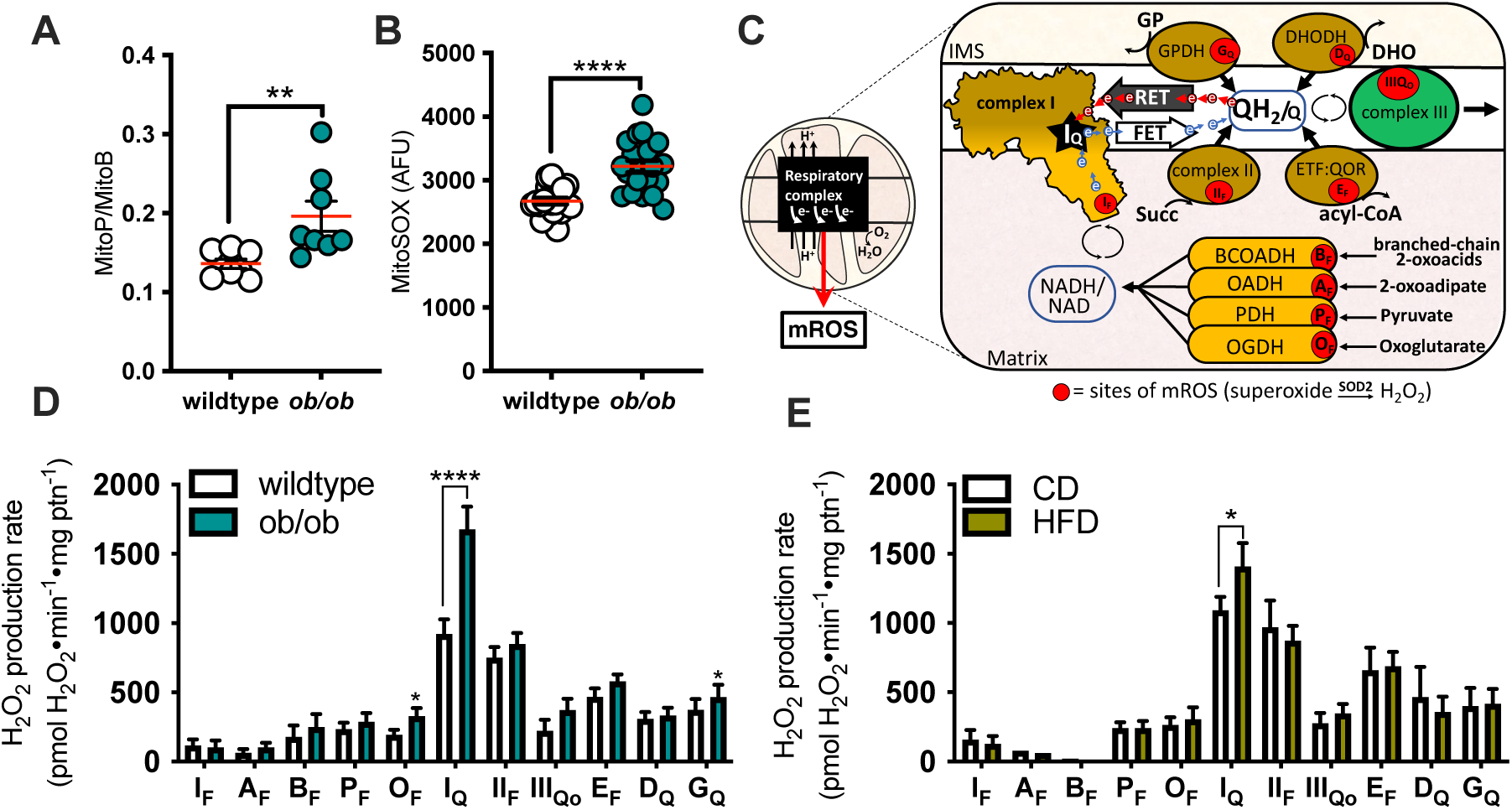
Reverse electron transport drives excess superoxide/H_2_O_2_ production in obese livers. (**A**) Quantification of H_2_O_2_ levels *in vivo* in the liver mitochondria of lean wildtype and obese leptin-deficient (ob/ob) mice. MitoP/MitoB ratio reflects H_2_O_2_ levels. Each dot represents a mouse (n=8). (**B**) Superoxide levels in primary hepatocytes measured by the oxidation of 1µM MitoSOX as a proxy. n= 281 (wildtype) and 138 (ob/ob) cells from 4 mice. AFU, arbitrary fluorescence units. (**C**) *Left*: Illustration of the general view of mROS generation, which ignores the individual sites that produce superoxide/H_2_O_2_. The lack of mechanistic information in this model hinders the identification of the precise source of superoxide in the mitochondria. *Right:* Illustration of the eleven sites from where electrons can directly reduce oxygen forming superoxide, which is then reduced to H_2_O_2_ by MnSOD (SOD2). These sites are represented as red circles. The NAD-linked sites are in the dehydrogenases of branched-chain 2-oxoacids (BCOADH, site B_F_), 2-oxoadipate (OADH, site A_F_), pyruvate (PDH, site P_F_) and oxoglutarate (OGDH, site O_F_). The CoQ-linked (Q) sites are in complex III (site III_Qo_), in the dehydrogenases of succinate (site II_F_), glycerol phosphate (GPDH, site G_Q_), dihydroorotate (DHODH, D_Q_) and in the electron transport flavoprotein: ubiquinone oxidoreductase (ETF:QOR, site E_F_). Complex I may have one or two sites that can transfer electrons to oxygen (*27*, *28*), the flavin site I_F_ and the Q-binding site I_Q_. Complex I generates superoxide during forward electron transport (FET) and reverse electron transport (RET). (**D**) Maximum capacity of superoxide/H_2_O_2_ production from liver-isolated mitochondria of wildtype and ob/ob mice (8-10 weeks of age, n=10-15 independent experiments). (**E**) Maximum capacity of superoxide/H_2_O_2_ production from liver-isolated mitochondria of wildtype mice fed a chow (CD) or a 60% high fat diet (HFD) for 17 weeks (n=6 independent experiments). The sites are described in panel (C). Values are means ± SEM. *, p<0.05; **, p<0.01; ***, p<0.0001 by Student’s *t* test.

Classically, complex I oxidizes NADH, transferring the electrons to ubiquinone (or coenzyme Q/CoQ) during forward electron transport (FET). However, complex I also catalyzes the reverse reaction, which is known as reverse electron transport (RET). During RET, electrons from the reduced CoQ pool (CoQH_2_) are forced back into complex I, driven by high protonmotive force (Δp), generating large amounts of superoxide (*14*, *21*). Although RET was once considered an *in vitro* artifact due to its unusual thermodynamic requirements, growing evidence shows that RET occurs *in vivo* and is physiologically relevant (*15*–*19*, *22*, *23*). However, how these individual sites and their activity states relate to metabolic health remains to be determined.

In this study, we systematically explored each of the 11 sites of mROS production in the liver tissues of lean and obese animals and determined that excess mROS in obesity is exclusively produced by RET from site I_Q_ in Complex I. Further, we determined that this unexpected biochemical alteration is due to ubiquinone deficiency and using multiple gain- and loss-of-function models *in vivo*, we demonstrated the importance of site I_Q_ RET for whole-body metabolism and its significance for human fatty liver diseases.

### Mitochondrial ROS generation is higher in the livers of obese mice

Current evidence for elevated mitochondrial reactive oxygen species (mROS) production in obesity has been generated exclusively using *ex vivo* and *in vitro* systems (*1*, *2*). To investigate hepatic mROS production in obesity *in vivo*, we first utilized the mitochondrial targeted ratiometric probe MitoB, which reacts with hydrogen peroxide (H_2_O_2_) to generate MitoP (*24*, *25*). The MitoP/MitoB ratio was greater in the livers of genetically obese (ob/ob) mice compared to lean controls, indicating higher levels of mitochondrial H_2_O_2_ in obesity *in vivo*, in the native tissue environment (**Fig. 1A**). We also confirmed that increased mROS in obesity was cell autonomous by treating primary hepatocytes with MitoSOX to measure superoxide levels. MitoSOX oxidation was elevated in primary hepatocytes from obese mice compared to lean controls (**Fig. 1B**) consistent with our earlier findings (*26*). Hence, obesity increases the oxidation status of liver mitochondria.

To determine the specific site(s) responsible for the excess mROS generation, we isolated mitochondria from livers of mice with either genetic or diet-induced obesity and their respective lean controls. To assess their maximum capacity of superoxide/H_2_O_2_ production we pharmacologically isolated each of the 11 sites and provided their appropriate substrate in the presence of oligomycin (**Fig. 1C-E**). The rate of superoxide/H_2_O_2_ were collectively measured as H_2_O_2_, which could be detected from all the sites. In isolated mitochondria from the liver of lean wildtype and chow fed (CD) mice, sites I_Q_ and II_F_ in complex I and II, respectively, had the highest capacity for mROS generation. Next, we focused on the only site that was exclusively upregulated in both models of obesity.

### Site I_Q_ in complex I generates excess mROS in the liver of obese mice via RET

Complex I is one of the major sources of mROS and its reaction is bidirectional (*29*), resulting in superoxide generation when the enzyme acts in both the forward and reverse electron transport (FET vs RET) (*30*, *31*) (**Fig. 1C and S1A**). However, multiple reports indicate that more superoxide is generated during RET than during FET (*14*, *30*–*32*). To measure the maximum capacity of mROS generation during FET from the flavin site of complex I (site I_F_), liver isolated mitochondria were incubated in the presence of malate, to generate NADH via the TCA cycle, aspartate and ATP, to suppress the mROS contribution from site O_F_ in oxoglutarate dehydrogenase (OGDH), and rotenone to prevent electrons from moving down the ETC (*33*) (**Fig. S1A**). We detected no difference in the rate of superoxide generation from site I_F_ via FET between mitochondria isolated from lean and obese livers (**Fig. 1D-E,** site I_F_). To induce mROS formation from site I_Q_ via RET, mitochondria were incubated with succinate, to increase CoQH_2_/CoQ ratio (CoQ redox state) and in the absence or presence of rotenone, a site I_Q_ specific inhibitor. RET was defined by the difference between these rates (**Fig. S1A and S1B**). Strikingly, site I_Q_ via RET was the only site differentially regulated in genetic and dietary models of obesity and its capacity was >80% and 40% elevated, respectively (**Fig. 1D-E,** site I_Q_). In addition to rotenone, S1QEL, piericidin A, and FCCP were also used to inhibit mROS production from site I_Q_ via RET (**Fig. S1A, S1C-E**). Consistently, mROS generation from site I_Q_ was higher in the liver isolated mitochondria of obese mice regardless the compound used to define RET (**Fig. S1F-G**). Notably, the increased superoxide/H_2_O_2_ production via RET in obesity appears specific to liver, as it was not observed in mitochondria from skeletal muscle of obese mice (**Fig. S1H**). Altogether, these data indicate that in isolated mitochondria from the livers of obese mice, excess mROS originates specifically from site I_Q_ in complex I via RET (**Fig. 1D-E**), indicating a site-specific dysregulation.

### Impaired CoQ synthesis in obesity drives mROS production via RET

To assess the mechanisms underlying increased RET in obesity, we systematically explored the thermodynamic forces that promote RET when elevated: i) the NADH redox state, ii) the magnitude of the mitochondrial Δp, and iii) the CoQ redox state. First, we monitored the NAD(P)H redox state in primary hepatocytes oxidizing pyruvate as an endogenous reporter of FET. Notably, the fraction of reduced NAD(P)H was significantly decreased in primary hepatocytes from obese mice, indicating decreased FET from complex I (**Fig. S2A-B**). Increased RET in obesity was also independent of changes in complex I or II activity, or overall changes in ETC protein levels (**Fig. S2C-F**). Next, we measured mitochondrial membrane potential as a proxy for Δp and found that succinate supported higher membrane potential in obese mitochondria, consistent with higher RET (**Fig. S2G-H**). This prompted us to investigate CoQ abundance and redox state directly. Interestingly, both the CoQ redox state and the fraction of reduced CoQ (CoQH_2_) were significantly increased in obese livers (**Fig. 2A-B**). This was associated with a significant decrease in total CoQ_9_ levels (**Fig. 2C**). Metabolomic analysis of *in situ* freeze-clamped liver tissues from lean and obese mice revealed no significant differences in the levels of metabolites that could potentially drive RET by directly feeding electrons into the CoQ pool, such as glycerol phosphate, dihydroorotate, acyl-carnitines, or succinate, which has been particularly shown to promote RET (*15*, *16*) (**Fig. S2I-M**). These observations rule out that the possibility that substrate accumulation promotes RET in obesity and support a model in which elevated CoQ redox state is the main driver of increased RET in the liver of obese mice.

**Fig. 2.**
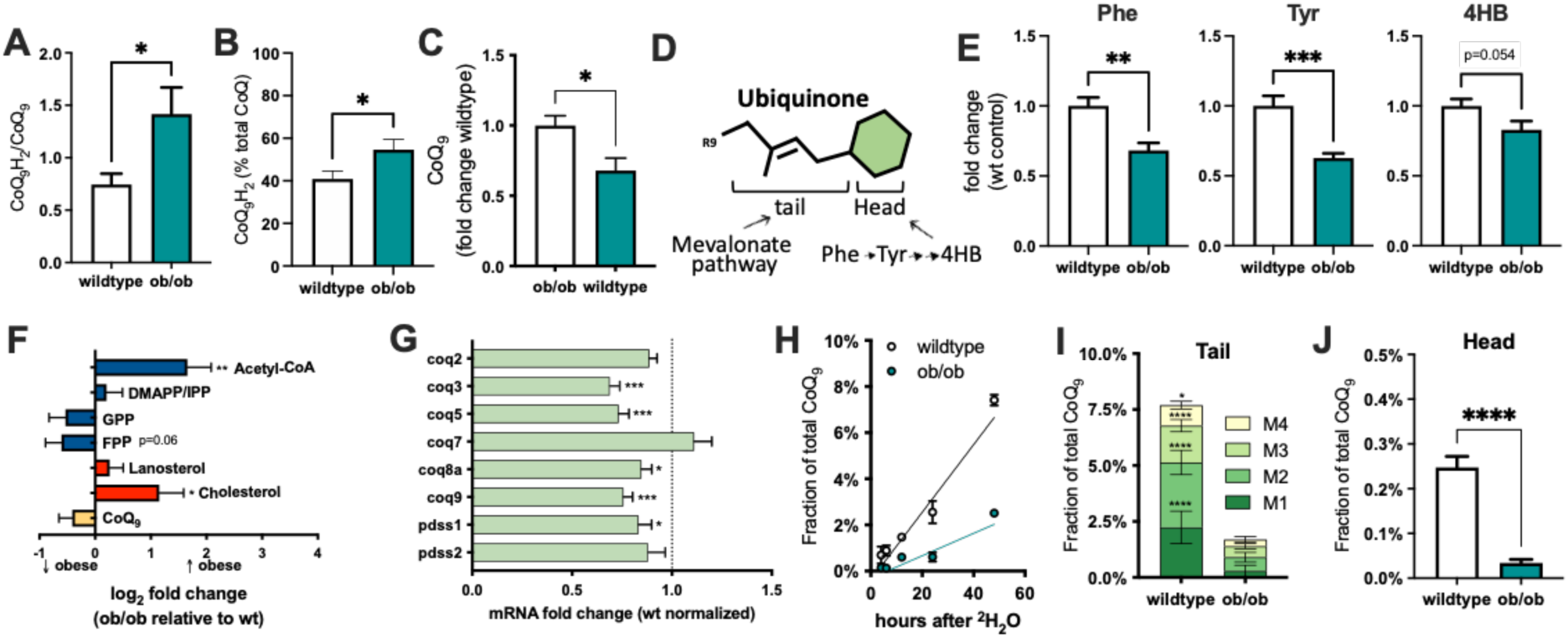
The CoQ redox state is increased in the livers of obese mice. (**A**) CoQ_9_ redox state (CoQH_2_/CoQ ratio) and (**B**) % of reduced CoQ (CoQH_2_) in the livers of wildtype and leptin-deficient ob/ob mice (8 weeks old, n=9 per group). (**C**) Total CoQ_9_ level in the liver of wildtype and ob/ob mice. (**D**) Illustration of ubiquinone (CoQ) chemical structure and precursors for the tail and head. The redox active head is derived from the metabolism of phenylalanine and tyrosine, while the isoprene tail, is derived from the mevalonate pathway. (**E**) Relative levels of the CoQ head precursors, phenylalanine (phe), tyrosine (tyr) and 4-hydroxibenzoate (4HB) in the livers of wildtype and ob/ob mice, 8-10 weeks of age. (**F**) Levels of metabolites in the mevalonate pathway (blue) and in the branches leading to the synthesis of cholesterol (red) and CoQ (yellow) in the livers of wildtype and ob/ob mice. Metabolite levels were normalized to wildtype livers and fold change was plotted as log_2_ values. (**G**) Relative expression levels of the genes in the CoQ biosynthetic pathway in the livers of wildtype and ob/ob mice. Expression levels were normalized to wildtype. (**H**) Kinetics of H^2^-enrichment in the CoQ_9_ pool in the livers of wildtype (n=10) and ob/ob mice (n=5). Mass enrichment in the CoQ_9_ isoprenoid tail (**I**) and head (**J**) in the livers of wildtype and ob/ob mice 24 hours after ^2^H_2_O administration in the drinking water (4% v/v) (n=6 per group). Values are means ± SEM. *, p<0.05; **, p<0.01; ***, p<0.001; ****, p<0.0001 by Student’s *t* test.

Next, we explored the mechanisms underlying lower hepatic CoQ_9_ levels in obesity utilizing metabolomics, gene expression and metabolic flux analysis. CoQ is comprised of a redox active quinone head attached to an isoprenoid tail of varying lengths (**Fig. 2D**) (*34*, *35*). The precursor for the head group is phenylalanine, which is converted to tyrosine and then 4-hydrobezoate (*34*), while the precursors for the tail are derived from the same mevalonate pathway as cholesterol (*34*, *35*). All three metabolites required for the head biosynthesis were decreased in the livers of obese mice (**Fig. 2E**). To investigate the potential nodes of regulation, we measured the levels of the metabolites in the mevalonate pathway and in the downstream branches leading to cholesterol and CoQ_9_ synthesis. Acetyl-CoA, an intermediate of fatty acid and glucose oxidation and a common precursor for *de novo* lipogenesis and the synthesis of sterols and non-sterols via the mevalonate pathway, was increased in the livers of obese mice (**Fig. 2F**). Conversely, farnesyl pyrophosphate (FPP), the last common metabolite in the synthesis of all products of the mevalonate pathway and direct CoQ_9_ precursor (*36*), was decreased in the livers of obese mice (**Fig. 2F**, FPP p=0.06). The gene expression profile of the mevalonate pathway did not show consistent alterations that would explain the changes in CoQ synthesis (**Fig. S2N**). On the other hand, genes directly responsible for CoQ biosynthesis including pdss1, which controls the lengthening of the CoQ tail, and coq3, coq5, coq8a and coq9, which are part of the mitochondrial complex Q and are necessary for CoQ head synthesis, were significantly decreased in livers of obese mice (**Fig. 2G**). These data suggest that in obesity, decreased CoQ biosynthetic processes contribute to the decreased CoQ levels.

To assess the *in vivo* kinetics of CoQ_9_ synthesis and the flux through the CoQ_9_ synthetic pathway in the liver, we provided lean and obese mice with ^2^H_2_O-supplemented drinking water for up to 48h and harvested livers for analysis. We observed a much slower rate of ^2^H incorporation into newly synthetized CoQ in the livers of obese mice (**Fig. 2H**), providing direct evidence for decreased CoQ synthesis *in vivo*. Next, we analyzed the flux through the synthetic pathways for the head and tail of CoQ_9_ at 24h after ^2^H_2_O administration. ^2^H incorporation in the CoQ_9_ isoprenoid tail and head was dramatically suppressed in the livers of obese mice (**Fig. 2I-J**). Taken together, the suppression of the CoQ biosynthetic program in the livers from obese mice drives a higher CoQH_2_/CoQ ratio, which promotes site-specific mROS via RET.

### Increased RET drives dysregulation of systemic glucose homeostasis

To test if excess mROS produced specifically via RET contributes to obesity-driven metabolic pathologies, we first pharmacologically induced complex I-mediated mROS formation using mitoparaquat (mitoPQ) (*25*, *37*), and tested whether this was sufficient to impair glucose metabolism. MitoPQ significantly enhanced mROS production via RET in a dose-dependent fashion when added to isolated mitochondria from the livers of lean mice (**Fig. 3A**). This effect was RET-specific, as mitoPQ had no effects on mROS production during FET or on the other mROS producing sites that feed electrons into the CoQ pool (**Fig. 3B and Fig. S3A**). MitoPQ-driven mROS production was sensitive to S1QEL, further supporting the conclusion that it originates from site I_Q_ (*38*) (**Fig. 3C and Fig. S3B-C**). Importantly, mitoPQ at the concentrations used here did not affect cellular bioenergetics, as treatment of primary mouse hepatocytes with different concentrations of mitoPQ for 60 minutes did not change basal or maximum oxygen consumption rates (**Fig. S3D**).

**Fig. 3.**
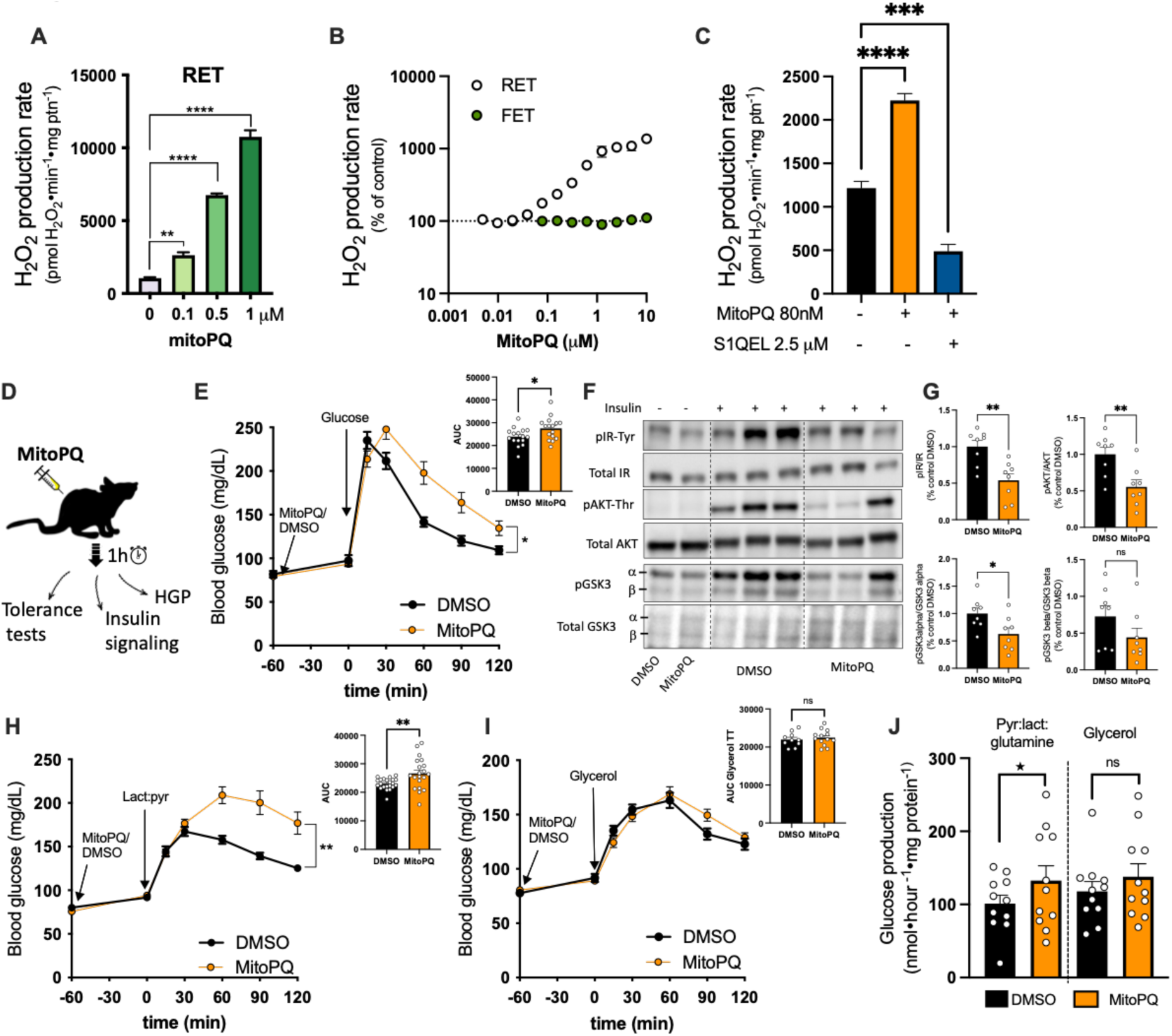
Pharmacological induction of RET in wildtype lean mice increases hepatic glucose production and impairs glucose homeostasis. (**A**) Mitoparaquat-induced (mitoPQ) superoxide/H_2_O_2_ generation via RET in mitochondria isolated from livers of wildtype (wt) lean mice is dose-dependent (n=3 independent mitochondrial isolations). (**B**) Effect of mitoPQ on the rate of superoxide/H_2_O_2_ production via RET or FET in wt isolated mitochondria. (**C**) MitoPQ-stimulated rate of superoxide/H_2_O_2_ production via RET is suppressed by 2.5µM S1QEL 2.2. (**D**) Illustration of acute mitoPQ treatment *in vivo*. (**E**) Blood glucose levels during intraperitoneal (i.p.) glucose tolerance test (0.5 g•kg^−1^) in 16h fasted wt mice treated with 4 nmol mitoPQ or DMSO vehicle as indicated, *inset* area under the curve. (**F**) Immunoblot and (**G**) quantification analysis of *in vivo* insulin signaling in total liver lysates of wt mice 1.5 h after treatment with 4 nmol mitoPQ or DMSO as vehicle (n=8 mice per group). (**H**) Blood glucose levels during i.p. lactate: pyruvate tolerance test (1.5 : 0.15 g•kg^−1^) in 16h fasted wt mice treated with 4 nmol mitoPQ or DMSO vehicle. *Inset* is area under the curve. (**I**) Blood glucose levels during i.p. glycerol tolerance test (1 g•kg^−1^) in 16h-fasted mice treated with 4 nmol mitoPQ or DMSO vehicle. *Inset* is area under the curve. (**J**) Glucose production (HGP) from primary hepatocyte utilizing 20 mM lactate, 2 mM pyruvate, and 2 mM glutamine or 20 mM glycerol as substrate. Hepatocytes were isolated from wt mice 1.5 h after mitoPQ or DMSO vehicle treatment (n=11 preparations per group). Values are means ± SEM. *, p<0.05; **, p<0.01; ***, p<0.001; ****, p<0.0001 by Student’s *t* test; back star, p<0.05 by paired t-test.

To assess the *in vivo* relevance of excess mROS production from site I_Q_ via RET, we administered mitoPQ intraperitoneally into healthy lean mice (**Fig. 3D**), which was well tolerated up to a dose of 4 nmol. Acute administration of mitoPQ impaired glucose tolerance in a dose-dependent manner (**Fig. 3E and Fig. S3E**) and decreased hepatic insulin signaling *in vivo* as measured by the decrease in the phosphorylated insulin receptor (IR), AKT and GSK3 alpha in the liver (**Fig. 3F-G**). The *in vivo* effects of mitoPQ-driven mROS via RET were liver-specific, since insulin signaling in the muscle and white adipose tissue were unchanged, and systemic insulin tolerance remained comparable to controls (**Fig. S3G-I**). Taken together, these data support a model in which excess mROS generated by RET *in vivo* is sufficient to cause hepatic insulin resistance, recapitulating the obese phenotype.

Gluconeogenesis utilizes a high proportion of substrates that are routed via the mitochondria such as pyruvate, lactate and glutamine (*39*). Other substrates, such as glycerol, provide gluconeogenic precursors relatively proximal in the pathway to the final product of glucose and downstream of mitochondrial function. We found that excess mROS production via RET induced remarkable increase in glucose excursion during a lactate:pyruvate tolerance test following mitoPQ administration (**Fig. 3H and Fig. S3F**), indicating a pronounced effect on lactate-supported gluconeogenesis. In contrast, glycerol-supported gluconeogenesis was not stimulated by mitoPQ (**Fig. 3I**), further confirming that the effect of mROS production via RET on hepatic glucose production strictly depends on mitochondrial metabolism. Furthermore, these effects were cell-autonomous, as primary hepatocytes isolated from mice injected with mitoPQ, but not vehicle, retained the capacity to produce more glucose when given substrates routed via mitochondria such as lactate, pyruvate, and glutamine, but not glycerol (**Fig. 3J**).

### Decreasing RET improves metabolic homeostasis in obesity

Next, we employed multiple genetic models of loss-of-function to test whether specifically targeting excess hepatic RET would have a positive metabolic outcome in obesity (**Fig. 4A**). In our first approach, we transfected two liver cell lines, Hepa 1-6 and AML-12, with *Ciona intestinalis* alternative oxidase (Aox). Aox is a cyanide-insensitive oxidase that provides an alternative route to oxidize excess CoQH_2_, decreasing the CoQH_2_/CoQ ratio and preventing mROS production via RET (**Fig. S4A-G**) (*40*–*42*). Consistently, Aox expression conferred cyanide-resistant oxygen consumption in these cells (**Fig. S4B-C**). Next, we isolated primary hepatocytes from obese mice and treated them with adenoviral (Ad) particles either expressing Aox or GFP. In these experiments, total H_2_O_2_ release was measured in the presence or absence of 5 µM S1QEL to assess the specific contribution of RET from site I_Q_ without blocking oxidative phosphorylation. H_2_O_2_ production from Ad.GFP-expressing hepatocytes was significantly more sensitive to S1QEL than that of Ad.Aox hepatocytes (**Fig. S4D**). The S1QEL-sensitive rate, referred to as RET (**Fig. S4E**), was also significantly lower in Aox-expressing hepatocytes from obese mice, consistent with the high CoQH_2_/CoQ ratio in obesity promoting excess mROS via RET. Furthermore, Aox expression was sufficient to improve insulin sensitivity in hepatocytes from ob/ob mice, as demonstrated by enhanced phosphorylation of insulin receptor (IR), AKT and GSK3 alpha (**Fig. S4F-G**).

**Fig. 4.**
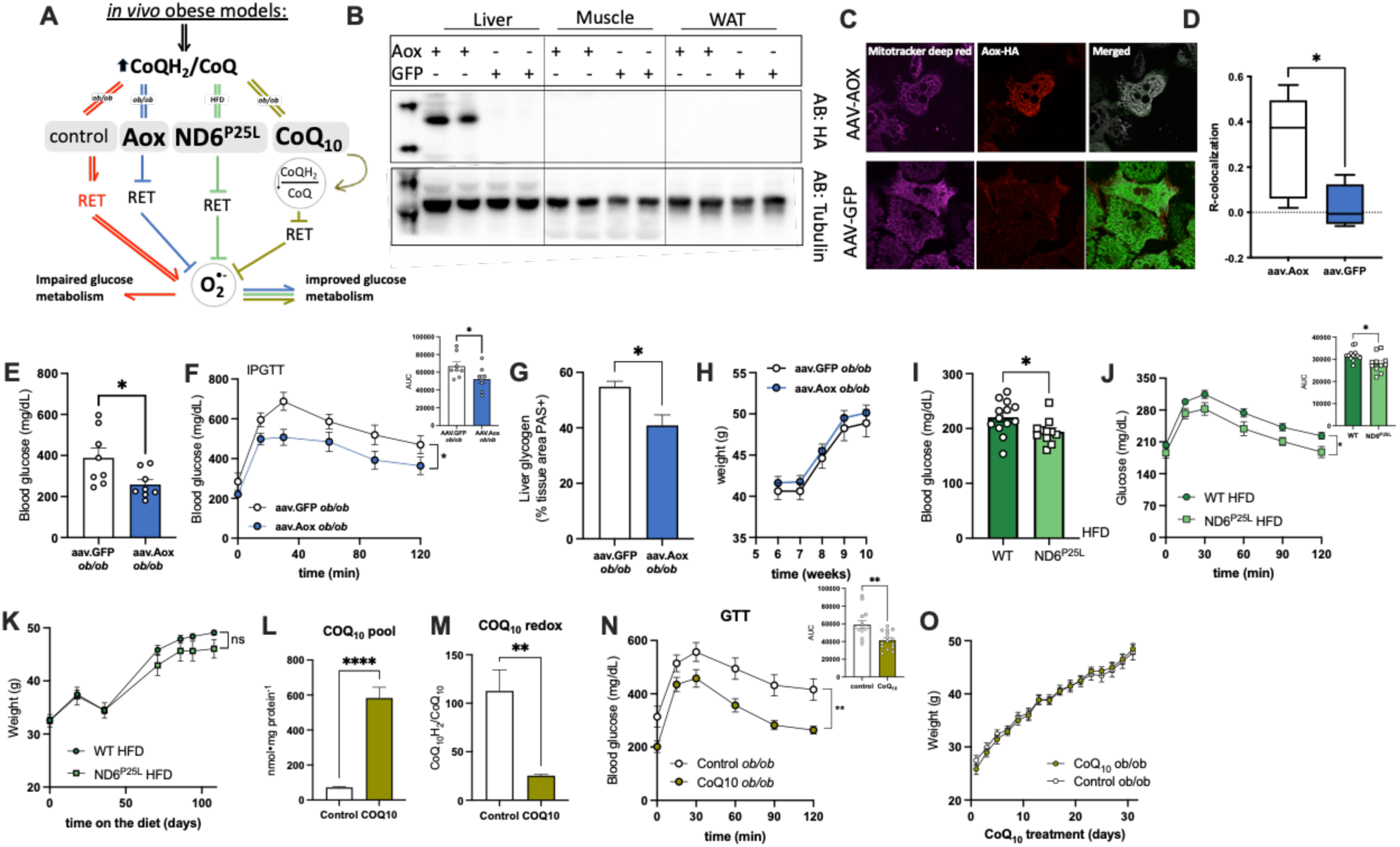
Suppressing RET *in vivo* improves metabolic outcomes in obese models. (**A**) Illustration to show that three independent loss-of-function models (Aox, ND6^P25L^ and CoQ_10_ supplementation) that suppress RET *in vivo* have improved glucose homeostasis. (**B**) Immunoblot confirming ectopic expression of *C. intestinalis* Aox in liver homogenates of ob/ob mice. (**C**) Representative confocal images of primary hepatocytes from aav.Aox or aav.GFP-expressing mice. Hepatocytes were loaded with MitoTracker deep red as a mitochondrial marker (purple, left panel) and HA-tagged-Aox was visualized with anti-HA antibody (red, middle panel). Superimposed images (white, right panel). (**D**) R-colocalization of Aox and MitoTracker deep red signals. (**E**) Six hour fasting blood glucose levels in ob/ob mice expressing aav.Aox or aav.GFP. (**F**) Blood glucose levels during i.p. glucose tolerance test (0.5 g•kg^−1^) in ob/ob mice expressing aav.Aox or aav.GFP. *Inset,* area under the curve. (**G**) Quantification of liver areas positive for PAS staining of glycogen in obese mice expressing aav.Aox or aav.GFP. (**H**) Body weights following aav.GFP or aav.Aox administration at 6 wks of age. (**I**) Six hour fasting blood glucose levels in wildtype and ND6^P25L^ mice fed high fat diet (HFD) for 15 weeks. (**J**) Blood glucose levels during i.p. glucose tolerance test (0.5 g•kg^−1^) in *ND6*^P25L^ and wildtype mice on HFD for 10 weeks. *Inset,* area under the curve. (**K**) Bodyweights of wildtype and ND6^P25L^ mice on HFD for 15 weeks. (**L**) CoQ_10_ levels and CoQ_10_ redox state (CoQH_2_/CoQ) (**M**) in the livers of obese mice treated with CoQ_10_ (10 mg•kg^−1^) or vehicle every other day for 4 weeks. (**N**) Blood glucose levels during i.p. glucose tolerance test (0.5 g•kg^−1^) in ob/ob mice treated with CoQ_10_ or vehicle. *Inset*, area under the curve. (**O**) Body weights of ob/ob mice treated with CoQ_10_ or vehicle. Values are means ± SEM. *, p<0.05; **, p<0.01; ****, p<0.0001 by Student’s *t* test or multiple comparisons analysis by 2-way ANOVA followed by Sidak’s post-hoc analysis.

Next, we tested whether preventing mROS production via RET *in vivo* could improve metabolic outcomes in obese mice. We utilized adeno-associated virus (AAV) to achieve hepatocyte-specific expression of Aox (**Fig. 4B and Fig. S4H**). Immunofluorescence and cellular fractionation demonstrated that Aox was specifically targeted to the mitochondria (**Fig. 4C-D and Fig. S4I**). In agreement with the role of RET in modulating glucose homeostasis (**Fig. 3**), hepatocytes isolated from obese mice expressing Aox displayed significantly lower glucose production in response to gluconeogenic substrates compared to mice expressing GFP (**Fig. S4J**). Consistent with these improvements at the cellular level, Aox-expressing mice exhibited significantly lower fasting blood glucose levels, improved intraperitoneal and oral glucose tolerance, and decreased liver glycogen content compared to GFP controls (**Fig. 4E-G and Fig. S4K**). Plasma insulin levels and insulin tolerance (**Fig. S4L-M**) were not changed, indicating that the systemic benefits of hepatic Aox expression were due to improved liver metabolism. Importantly, these metabolic improvements were independent of any changes in weight gain (**Fig 4H**), body composition, energy expenditure, respiratory exchange ratio (RER), or liver metabolites in Aox-expressing animals compared GFP control mice (**Fig. S4N-R**).

As a final additional test for the role of RET-mediated mROS production in promoting metabolic dysfunction during obesity, we utilized a mouse model that is incapable of generating mROS via RET (*43*, *44*). The mitochondrial DNA mutation G13997A causes a proline to leucine substitution in position 25 of the *ND6* gene (*ND6*^P25L^), which encodes the NADH dehydrogenase subunit 6 (ND6) of complex I. After 16 weeks on a high fat diet (HFD), *ND6*^P25L^ mice exhibited lower fasting blood glucose levels and were significantly more glucose tolerant than wildtype mice (**Fig. 4I-J**). This difference was not observed in chow fed mice (**Fig. S5A-C**) and was independent of changes in body weight (**Fig. 4K**). These results show that mROS production via RET has a causative role in disturbing glucose homeostasis in obesity.

### CoQ redox state in patients with hepatic steatosis favors mROS via RET

The prospect that CoQ deficiency drives mROS via RET and contributes to the pathological consequences of obesity raises the possibility that CoQ supplementation, if properly delivered to liver, could prevent or ameliorate some obesity-related pathologies. To investigate this question, we first treated young ob/ob mice with a new CoQ_10_ emulsion formulation via intraperitoneal injection. CoQ_10_ treatment (10mg/kg) every other day for 4 weeks significantly increased CoQ_10_ levels in the livers of obese mice and lowered the CoQH_2_/CoQ ratio (**Fig. 4L-M**). These changes were sufficient to improve whole body glucose tolerance (**Fig. 4N**) and insulin sensitivity in obese mice (**Fig. S5D**). CoQ_10_ supplementation had no effect on glucose tolerance in lean mice (**Fig. S5E**) or body weight in either lean or obese mice (**Fig. 4O and S5F**).

To investigate the relevance of the CoQ synthetic pathway and redox state in obese humans with liver disease, we analyzed samples from two cohorts of obese individuals with various degrees of nonalcoholic fatty liver disease (NAFLD, steatosis grade S0-S3) (**Tables S3-S4**). In the first cohort, we measured the expression of genes responsible for CoQ synthesis, which revealed that transcripts from the CoQ synthetic pathway were significantly lower in patients with steatosis, relative to healthy controls (**Fig. 5A**). Indeed, the expression levels of coq2, coq3, coq6, coq7 and pdss1 were consistently decreased in all steatosis stages.

**Fig. 5.**
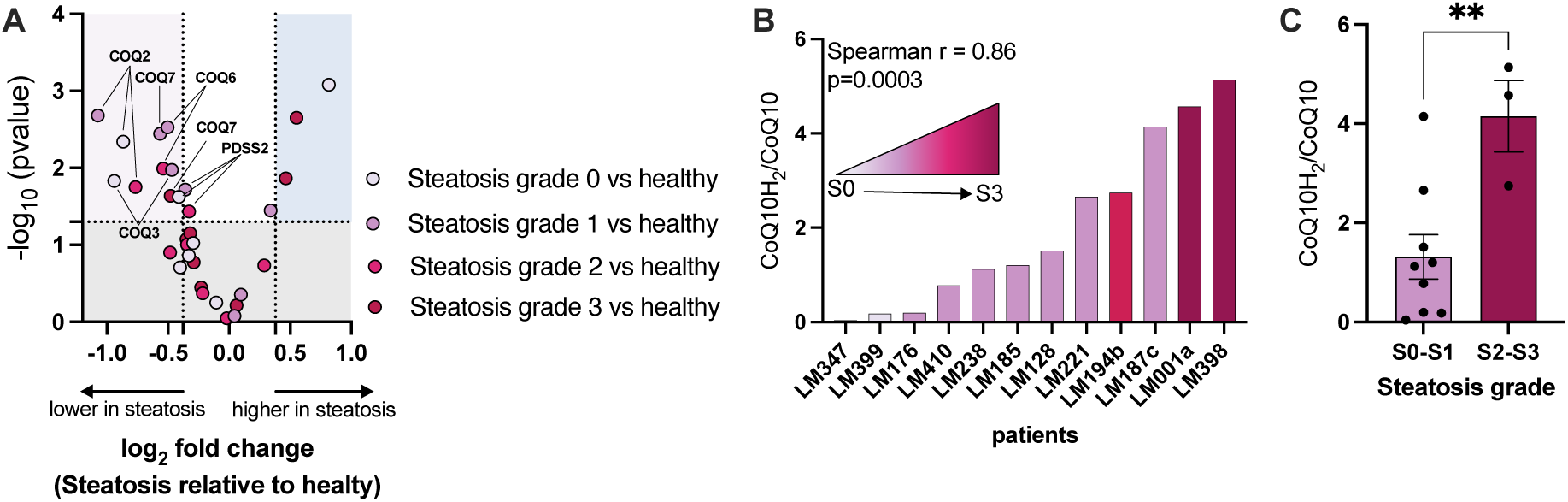
CoQ - RET axis in hepatic steatosis in humans. (**A**) Transcript levels of the enzymes from the CoQ biosynthetic pathway in liver biopsy samples from patients at different stages of steatosis compared to healthy subjects. (**B**) Correlation between CoQ_10_ redox state and the stages of steatosis in liver biopsies from human patients. (**C**) Comparison of CoQ_10_ redox state between patients with low- and high-grade steatosis. Values are means ± SEM. **, p<0.005 by Student’s *t* test.

Next, we measured the CoQH_2_/CoQ ratio in hepatic tissues in a second cohort of obese individuals with NAFLD. To reliably capture the *in vivo* CoQH_2_/CoQ ratio, tissue biopsies (∼2 mg) were immediately placed in liquid nitrogen after surgery. In these samples, we found a strong positive correlation between the CoQ redox state and the steatosis grade (Spearman r=0.8, p=0.0003) (**Fig. 5B**). Additionally, when stratified by steatosis grade, individuals with more severe steatosis (S2-S3) had a significantly higher CoQH_2_/CoQ ratio than those with lower grades (S0-S1) (**Fig. 5C**). These data suggest that excess mROS production via RET can also occur in human livers from obese individuals, and this may be due to lower CoQ synthesis and a higher CoQ redox state. Finally, given that CoQ deficiency may drive metabolic disease, pharmacological interventions that disrupt CoQ synthesis may worsen disease progression, whereas interventions that replenish CoQ may be promising therapeutics for metabolic disease.

## Discussion

Our data provide direct evidence that excess hepatic mROS is generated from site I_Q_ in complex I via RET in obesity. We describe a previously unrecognized mechanism in which a defect in CoQ biosynthesis in the liver mitochondria of obese mice overwhelms the CoQ pool, increasing the CoQH_2_/CoQ ratio. This high CoQ redox state creates the necessary thermodynamic force to drive electrons in reverse to complex I, generating excess mROS. Here, we utilized a combination of genetic models and pharmacological interventions to demonstrate that excess mROS production via RET at complex I is sufficient to impair metabolism in lean mice, while preventing hepatic RET in obesity is sufficient to improve glucose and lipid homeostasis.

Although RET has been observed for many years in isolated mitochondria, the understanding of its pathophysiological significance is much more recent (*42*, *45*). RET has important homeostatic roles (*17*, *18*), but is also linked to pathological superoxide production, such as in ischemia-reperfusion injury (*15*), and mediates inflammation and susceptibility to infection (*16*, *22*, *23*). Under these conditions, RET is triggered by a rapid oxidation of accumulated of succinate (*15*, *16*, *19*). Importantly, the mechanism described in this study is different in several ways. First, it is independent of substrate accumulation, a feature that has not been described before. Second, we find that RET activation in obesity relies primarily on the defect in CoQ synthesis. Finally, RET in this context is connected to local and systemic metabolic homeostasis.

We found that the CoQ biosynthetic pathway is decreased in the livers of obese individuals, and that the CoQ redox state strongly correlates with the severity of hepatic steatosis. These observations support that the mechanism described here may be relevant for human biology and the pathogenesis of fatty liver disease. Furthermore, our findings may promote re-imagining of the approaches for CoQ supplementation. The electrons from all mitochondrial inner membrane dehydrogenases converge into CoQ, thus the CoQ redox state is a metabolic sensor, as an elevated CoQH_2_/CoQ ratio flags higher electron input over output (*46*). In fatty liver diseases and states of insulin resistance, fatty acid oxidation is increased (*47*–*49*), potentially contributing to the higher CoQ redox state that favors RET. CoQ is also a lipid-soluble antioxidant found in other cellular membranes and in circulation (*50*), therefore carrying the ability to influence multiple functional domains of the cells. CoQ_10_ levels in circulation are often decreased in patients prescribed with statins to control their cholesterol levels (*51*). Statin use is also associated with an increased risk of developing type 2 diabetes (*52*–*54*), although the mechanism is still unclear. Our findings suggest that pharmacological interventions that disrupt CoQ synthesis may worsen metabolic diseases, while therapeutics to replenish CoQ, including during statin use, may be beneficial. In fact, CoQ supplementation has ascended in popularity in recent years, especially in patients prescribed with statins. However, these approaches have failed to show beneficial effects on glucose and lipid homeostasis (*55*, *56*), perhaps by missing the optimum dose or by lacking the ability to deliver CoQ to the target tissue. Here, we successfully delivered CoQ to the liver by using a different CoQ formulation and showed that it is effective in correcting glucose metabolism.

In conclusion, our finding that excess hepatic mROS generation in obesity is site-specific opens new perspectives for the development of therapeutic strategies focused on either decreasing RET, increasing CoQ levels, or both.

## Acknowledgements

We thank K. Prentice for the critical review of the paper; L. Lambourne for help and advice on statistical analysis; K. Langston for the help with qPCR; We are in debt to N. Snyder, E. Cagampan, N. Min, L. Greene, S. Karzhevsky for their technical assistance; D. Wallace for generously sharing the ND6-P25L mouse line; E. Dufour and P. Rustin for providing the initial Aox plasmid and antibody. C. Vidoudez and S. Trauger for helping with mitoB/P and CoQ quantifications; Z. Kahn and the HSPH animal facility staff for animal husbandry; and M. Rodriguez for laboratory maintenance.

## Author Contributions

Conceptualization: R.L.S.G., G.S.H.

Methodology: R.L.S.G., G.P., S.C.B., X.F., G.Y.L., K.I.,

Software: G.P.

Validation: R.L.S.G., Z.B.W., X.F., A.P.A., J.S., C.R.

Formal Analysis: R.L.S.G.

Investigation: R.L.S.G.

Visualization: R.L.S.G., G.S.H.

Resources: G.S.H., R.L.S.G., S.C.B., S.T.H, I.G., M.C.L.

Writing – original draft: R.L.S.G.,

Writing – review & editing: R.L.S.G, G.S.H, A.P.A, G.P. S.C.B., K.I., J.S., G.Y.L.

Funding acquisition: R.L.S.G., G.S.H.

Project Administration: R.L.S.G., G.S.H.

Supervision: R.L.S.G., G.S.H.

## Data and materials availability

All data necessary to evaluate the conclusions presented in this manuscript are present in the paper and/or the Supplementary Materials. The microarray data is available on the GEO repository (accession numbers GSE139602 and GSE224645).

## Methods

### General animal care, study design and animal models

Mouse husbandry and experiments were in compliance and approved by the Harvard Medical Area Standing Committee on Animals. Mice were maintained from 4-20 weeks on a 12-hour-light /12-hour-dark cycle, with lights on at 7:00 am, in the Harvard T.H. Chan School of Public Health pathogen-free barrier facility. Mice had free access to water and standard laboratory chow diet (PicoLab Mouse Diet #5058, LabDiet), unless otherwise indicated.

### Animal models

The obese models used in this study were the leptin-deficient B6.Cg-Lep^ob^/J (ob/ob) mice on a C57BL/6J genetic background, age and sex-matched ob/+ heterozygous (ob/ob and ob/+ Jackson Laboratories, stock no. 000632, Bar Harbor, ME, USA) were used as controls. For the diet-induced obesity model, C57BL/6J mice (Jackson Laboratories, stock no. 000664) were placed on high fat diet (#D12492i 60% kcal from fat, Research Diets) for 18 weeks starting at 6 weeks of age. Control mice were placed on a low-fat chow diet (PicoLab Mouse Diet #5053, LabDiet). The mouse strain ND6-P25L, which carries the single point mutation in the mitochondrially encoded ND6 gene, was a kind gift from Professor Douglas Wallace, Children’s Hospital of Philadelphia, Un. of Pennsylvania. As female carrying the mutation were backcrossed over several generations onto C57BL/6J background we used wildtype mice as controls. For that, females with the ND6 mutation and C57BL/6J wildtype females were crossed with wildtype male mice to generate the cohort used in this study. Wildtype and ND6-P25L mice were placed on control or high fat diet for 15 weeks starting at 6-7 weeks of age.

### Mitochondria isolation

Mouse livers were minced with a razor blade in a glass petri dish containing ice-cold STE buffer [250mM sucrose, 5mM Tris-HCl, 2 mM EGTA, pH 7.4 at 4°C]. Liver pieces were rinsed in ice-cold STE and homogenized in a glass homogenizer with a loose-fit glass pestle using six to nine strokes. The homogenate was filtered through a double layer of gauze and ∼ 15 mL of STE was added to the filtrate, which was then centrifuged at 1,000 g for 3 min. The supernatant was centrifuged at 10,000 g for 10 min to pellet the mitochondria and the pellet was washed twice in STE medium. The final crude mitochondrial pellet was resuspended in 1 mL STE and underwent a further purification step by Percoll density centrifugation using a discontinuous density gradient of 2 mL 80%, 4.71 mL 52% and 4.71 mL 26% Percoll in 2 mL 4x STE to a final Percoll density gradient of 1.117, 1.08, 1.047 g/mL, respectively, at 15,500 rpm for 45 min, at 4°C, using a 40Ti swing bucket in an XL-90 Beckman centrifuge. The mitochondria fraction was collected between the 52%-26% interfaces and washed once in STE buffer at 10,000 g for 10 min to obtain purified mitochondria. Mitochondria were resuspended in 1mL STE medium and immediately assayed.

Mouse skeletal muscle mitochondria were isolated as previously described (*57*). Briefly, skeletal muscle was dissected from the entire hind limbs of wildtype and ob/ob mice and placed in 4°C Chappell-Perry buffer (CP1, 0.1 M KCl, 50 mM Tris, 2 mM EGTA, pH 7.4) and CP2 (CP1 supplemented with 0.5 % w/v fatty acid-free BSA, 2 mM MgCl2, 1 mM ATP and 250 U•0.1 ml^−1^ subtilisin protease type VIII, pH 7.4) and mitochondria were isolated as described previously (*58*), resuspended in CP1 medium and immediately assayed. Protein concentration was determined using bicinchoninic acid assay.

### Superoxide/H_2_O_2_ measurements

Rates of superoxide/H_2_O_2_ production were collectively measured as rates H_2_O_2_ production as endogenous and exogenous superoxide dismutase (SOD) convert two superoxide molecules to one H_2_O_2_. H_2_O_2_ generation in purified liver mitochondria (0.1 mg•ml^−1^) was measured during state 4 (oligomycin 2.5 μg•ml^−1^) in KHE assay buffer [120 mM KCl, 5mM HEPES, 1 mM EGTA] supplemented with 2 mM MgCl_2_, 10 mM KH_2_PO_4_, 0.3 % w/v fatty acid-free BSA, pH 7.1 at 37°C. The detection system consisted of 0.2 U•ml^−1^ horseradish peroxidase (HRP), 50 μM Amplex UltraRed (Molecular Probes), 100 μM PMSF, 24 U•ml^−1^ SOD. In liver-isolated mitochondria it is mandatory to add PMSF to inhibit the non-specific conversion of Amplex UltraRed to resorufin (*59*). Skeletal muscle mitochondria (0.3 mg•ml^−1^) were resuspended in KHE medium supplemented with 1mM MgCl_2_, 5 mM KH_2_PO_4_, 0.3 % w/v fatty acid-free BSA, pH 7.1 at 37°C in the presence of oligomycin 2.5 μg•ml^−1^. Similarly, the detection system was 5 U•ml^−1^ HRP, 50 μM Amplex UltraRed, 25 U•ml^−1^ SOD. Fluorescence was monitored using a microplate reader (SpectraMax Paradigm, Molecular Devices) at Λ_excitation_= 605 nm, Λ_emission_= 640 nm after wavelength optimization. Fluorescence was calibrated using H_2_O_2_ standards after each run under the exact conditions of the assay.

**Table S1.**
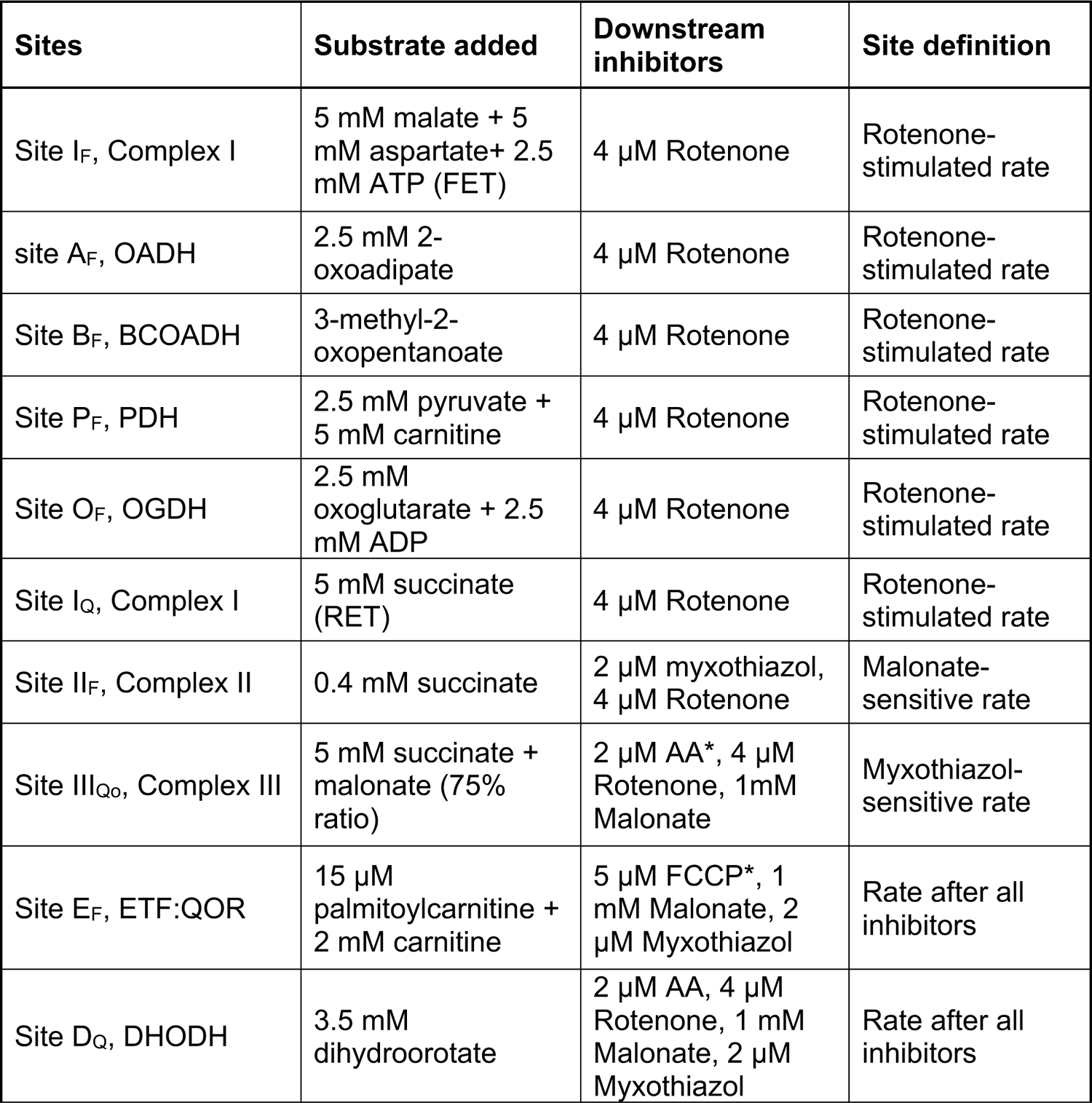
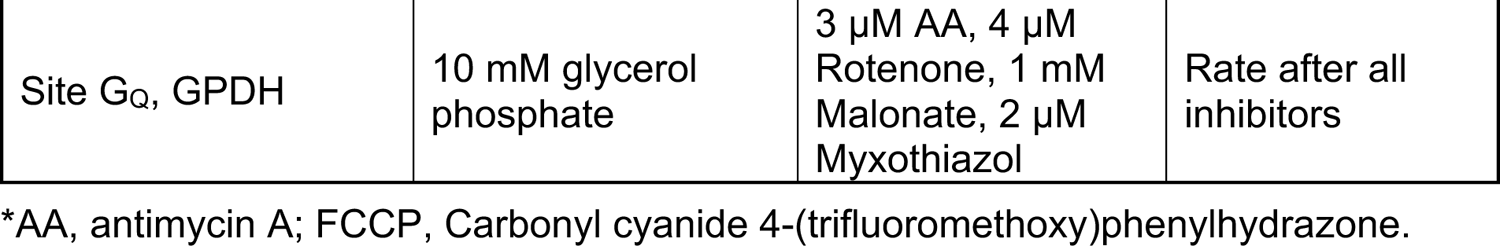
Substrates and inhibitors used to measure site-specific superoxide production in isolated mitochondria.

| Sites | Substrate added | Downstream inhibitors | Site definition |
| --- | --- | --- | --- |
| Site I<sub>F</sub>, Complex I | 5 mM malate + 5 mM aspartate+ 2.5 mM ATP (FET) | 4 $\mu\text{M}$ Rotenone | Rotenone-stimulated rate |
| site A<sub>F</sub>, OADH | 2.5 mM 2-oxoadipate | 4 $\mu\text{M}$ Rotenone | Rotenone-stimulated rate |
| Site B<sub>F</sub>, BCOADH | 3-methyl-2-oxopentanoate | 4 $\mu\text{M}$ Rotenone | Rotenone-stimulated rate |
| Site P<sub>F</sub>, PDH | 2.5 mM pyruvate + 5 mM carnitine | 4 $\mu\text{M}$ Rotenone | Rotenone-stimulated rate |
| Site O<sub>F</sub>, OGDH | 2.5 mM oxoglutarate + 2.5 mM ADP | 4 $\mu\text{M}$ Rotenone | Rotenone-stimulated rate |
| Site I<sub>Q</sub>, Complex I | 5 mM succinate (RET) | 4 $\mu\text{M}$ Rotenone | Rotenone-stimulated rate |
| Site II<sub>F</sub>, Complex II | 0.4 mM succinate | 2 $\mu\text{M}$ myxothiazol, 4 $\mu\text{M}$ Rotenone | Malonate-sensitive rate |
| Site III<sub>QO</sub>, Complex III | 5 mM succinate + malonate (75% ratio) | 2 $\mu\text{M}$ AA*, 4 $\mu\text{M}$ Rotenone, 1mM Malonate | Myxothiazol-sensitive rate |
| Site E<sub>F</sub>, ETF:QOR | 15 $\mu\text{M}$ palmitoylcarnitine + 2 mM carnitine | 5 $\mu\text{M}$ FCCP*, 1 mM Malonate, 2 $\mu\text{M}$ Myxothiazol | Rate after all inhibitors |
| Site D<sub>Q</sub>, DHODH | 3.5 mM dihydroorotate | 2 $\mu\text{M}$ AA, 4 $\mu\text{M}$ Rotenone, 1 mM Malonate, 2 $\mu\text{M}$ Myxothiazol | Rate after all inhibitors |
| Site G<sub>Q</sub>, GPDH | 10 mM glycerol phosphate | 3 μM AA, 4 μM Rotenone, 1 mM Malonate, 2 μM Myxothiazol | Rate after all inhibitors |

\*AA, antimycin A; FCCP, Carbonyl cyanide 4-(trifluoromethoxy)phenylhydrazone.

### Mitochondrial membrane potential with Safranine O, complex I and complex II activities

Mitochondrial membrane potential was estimated in purified liver mitochondria (0.1 mg•ml^−1^) in the presence of 2.5 μg•ml^−1^ oligomycin during succinate oxidation in KHE assay buffer [120 mM KCl, 5 mM HEPES, 1 mM EGTA] supplemented with 2 mM MgCl_2_, 10 mM KH_2_PO_4_, 0.3 % w/v fatty acid-free BSA, pH 7.1 at 37°C. The detection system consisted of 2 µM Safraine O and fluorescence was monitored using a microplate reader (SpectraMax Paradigm, Molecular Devices) at Λ_excitation_= 495 nm, Λ_emission_= 587 nm. Freeze-thawed mitochondria (2.5 mg•ml^−1^) were incubated in 250 mM sucrose, 0.2 mM EDTA, 50mM Tris-HCl (pH 7.0) and 5 µM cytochrome c. NADH oxidation (150 µM) was monitored at 340 nm. Complex I activity was expressed as the rotenone-sensitive rates of NADH oxidation. Complex II activity was measured using MitoCheck Complex II Activity Assay Kit (Cayman, USA).

### *In vivo* H_2_O_2_ levels with MitoB

The levels of H_2_O_2_ *in vivo* in the liver mitochondria were determined according to (*24*). Lean and ob/ob 7-week-old mice were injected with 0.8 µmol•kg^−1^ MitoB (Cayman Chemical) via the tail vein. Six hours after injection livers were collected and immediately frozen. MitoB and MitoP were extracted from 100 mg of tissue, as previously described, and labelled internal standards d-_15_ MitoB and d-_15_ MitoP were added to each sample (*24*). The samples were analyzed using multiple reaction monitoring (MRM) on an Agilent 6460 LC-MS/MS triple quad mass spectrometer. LC conditions were slightly modified from Cocheme 2012. Five microliters of each sample were injected on a PhenylHexyl column (Dikma Platisil, 5um, 150×4.6mm). Mobile phases were A: water with 0.1% formic acid. B: acetonitrile with 0.1% formic acid. The gradient was 5% B for 2 min, to 25% B in 1min, to 60% B in 3.5min and then to 100% B in 3min. After 6.5 min at 100% B the column was re-equilibrated at 5% B for 3min. The flow rate was maintained at 0.3mL/min. MitoP and MitoB concentrations were determined using calibration curves from pure standards prepared similarly to the samples, using the d-_15_ labelled compounds as internal standards.

### CoQ isolation and redox

CoQ was extracted from clamp frozen livers of lean and obese mice to ensure that the CoQH_2_/CoQ ratio (CoQ redox state) accurately reflected the *in vivo* state. After cervical dislocation the liver was immediately dissected and kept at −80°C until the experiment. 5-6 mg of frozen tissue was homogenized in 250 μL ice-cold acidified methanol (0.1% HCl) and 250 μL water washed with hexanes in a bullet blender (TissueLyser II, Qiagen) with 5 mm stainless steel beads (Qiagen) at 30 Hz for 3 min at 4°C. The samples were spiked with 200 pmol CoQ_8_ as internal standard and centrifuged at 17000 g for 5 min. The upper layer containing CoQ and was dried under argon and then resuspended in methanol containing 2 mM ammonium formate and analyzed within 24 hours. The same protocol was used to extract CoQ from human liver biopsies, however the average starting material was 2 mg. The reduced CoQ standards were prepared according to published methodology (*60*). Samples were analyzed in a Thermo QExactive Plus mass spectrometer coupled to an Ultimate 3000 LC (Thermofisher). Five microliters of sample were injected onto an Eclipse Plus C18 column (Agilent, 2.1×50mm). Mobile phases were A: methanol, 5mM ammonium formate. B: isopropanol. The gradient was as follows: 9 min at 20% B, then to 100% B in 0.1 min. After 4 min at 100% B the column was re-equilibrated to 20% B for 3 min. The flow rate was 0.3 mL/min for the first 9 min and then was switched to 0.4 mL/min. Compounds were quantified using the signal of the high resolution mass of the [M+NH_4_]+ adducts. Quantification was done using the standard curves and CoQ_8_ as the internal standard.

### Measurement of CoQ enrichment by LC-MS/MS

CoQ_9_ and CoQ_10_ enrichment were measured using a modified reverse-phase liquid chromatography-electrospray ionization-tandem mass spectrometry (LC-MS) method (*60*). Approximately 20 mg of frozen tissue was homogenized in 1 mL methanol. The solution was transferred to a larger tube, and the tube was washed with another 1 mL methanol which was combined with the solution. Hexane (10 mL) was added and the samples were vortexed and then centrifuged for 5 min at 1635 *g*. The supernatants were then removed and dried under N_2_. The dried extracts were reconstituted in 180 μL of methanol/isopropanol (80:20, vol/vol) with 5 mM ammonium acetate before analysis by LC–MS.

LC was performed using a reverse phase Luna C8 column (Phenomenex, 150 × 2.1 mm, 5 µm). The mobile phase consisted of methanol with 5 mM ammonium acetate (eluent A) and isopropanol (eluent B). The gradient consisted of 20% B for 8 min, and was increased to 100% B over 0.5 min. After 10 minutes the mobile phase was switched to 20% B over 0.5 min and maintained for 7 min. The analytes were detected by ESI-MS/MS using an API 3200 triple-quadrupole LC-MS/MS system equipped with an ESI Turbo Ion Spray interface, operated in the positive ion mode (AB Sciex, Framingham, MA, USA). The ion source parameters were set as follows: curtain gas, 20 psi; ion spray voltage, 5000 V; ion source temperature, 350°C; nebulizing and drying gas, 30 and 40 psi; declustering potential, 20 V; and collision energy, 50 V. Triple-quadrupole scans were acquired in the multiple reaction monitoring (MRM) mode with Q1 and Q3 set at “unit” resolution. MRM transitions for m0, m1, m2, m3 and m4 mass isotopomers of deuterated CoQ_10_ and CoQ_9_ are summarized in Table 2. Resulting isotopomers were corrected for natural abundance.

**Table S2.** MRM transition of CoQ_10_ and CoQ_9._

| Compound | Isotopomers | Q1 m/z | Q3 m/z |
| --- | --- | --- | --- |
| CoQ<sub>10</sub>H<sub>2</sub> (Reduced) | m0 | 882.8 | 197.1 |
|  | m1 | 883.8 | 197.1 |
|  | m2 | 884.8 | 197.1 |
|  | m2 | 884.8 | 199.1 |
|  | m3 | 885.8 | 197.1 |
| CoQ<sub>9</sub>H<sub>2</sub> (Reduced) | m0 | 814.8 | 197.1 |
|  | m1 | 815.8 | 197.1 |
|  | m2 | 816.8 | 197.1 |
|  | m2 | 816.8 | 199.1 |
|  | m3 | 817.8 | 197.1 |
|  | m4 | 818.8 | 197.1 |
| CoQ<sub>9</sub> (oxidized) | m0 | 812.8 | 197.1 |
|  | m1 | 813.8 | 197.1 |
|  | m2 | 814.8 | 197.1 |
|  | m2 | 814.8 | 199.1 |
|  | m3 | 815.8 | 197.1 |
|  | m4 | 816.8 | 197.1 |
| CoQ<sub>10</sub> (oxidized) | Not detected |  |  |

### ^2^H_2_O treatment

Wildtype and ob/ob mice were administered deuterated water (^2^H_2_O) for 48 hours. Initially, mice were injected with a bolus of saline solution 0.9% (w/v) prepared in ^2^H_2_O at a dose 27µL•gram body weight ^−1^. Mice were then offered ^2^H_2_O (4% v/v) in their drinking water. Liver tissues were collected at 4, 6, 12, 24 and 48h and prepared for CoQ_9_ extraction and measurement of H^2^-enrichement in the CoQ tail and head.

### Cell culture

Hepa 1-6 cells, a mouse liver hepatoma cell line, were cultured in DMEM with 10% CCS. AML12 cells, a mouse normal hepatocyte cell line, were cultured in DMEM:F12 Medium with 10% FBS, 1x STI, 40ng/mL dexamethasone. All cells were cultured at 37°C in a humidified incubator that was maintained at a CO_2_ level of 10%.

### Primary hepatocyte isolation

Mice were anesthetized with 200 mg•kg^−1^ ketamine and 20mg•kg^−1^ xylazine and livers were then perfused with 50 mL of buffer I [11 mM glucose; 200 μM EGTA; 1.17 mM MgSO_4_ heptahydrate; 1.19 mM KH_2_PO_4_; 118 mM NaCl; 4.7 mM KCl; 25 mM NaHCO_3_, pH 7.32] via the portal vein until the liver turned pale, using an osmotic pump set to an infusion rate of ∼4 mL•min^−1^. The rate was then gradually increased to ∼7 mL•min^−1^. After ∼ 5 min, when the entire buffer I had been infused, 50 mL of buffer II [11 mM glucose, 2.55 mM CaCl_2_, 1.17 mM MgSO_4_ heptahydrate, 1.19 mM KH_2_PO_4_, 118 mM NaCl, 4.7 mM KCl, 25 mM NaHCO_3_, 7.2 mg/mL fatty acid-free BSA, 0.18 mg•mL^−1^ of Type IV Collagenase (Worthington Biochem Catalog: LS004188], was infused. BSA and collagenase were added to buffer II immediately before infusion. The buffers were kept at ∼37°C during the entire perfusion. Next, the liver was excised, and primary hepatocytes were carefully released and sedimented at 500 rpm for 2 min, washed twice in DMEM with 10% CCS and suspended in Williams E medium supplemented with 5 % CCS and 1 mM glutamine (Invitrogen, CA). To separate live from dead cells, the hepatocytes were layered on a 30% Percoll gradient and centrifuged at ∼1500 rpm for 15 min. The healthy cells were recovered at the bottom of the tube and plated in Williams E medium and kept overnight until experimentation.

### Hepatocyte oxygen consumption

Hepatocyte oxygen consumption rates (OCR) were monitored using an XF-24 and XF-96 extracellular flux analyzer (Seahorse Bioscience; Agilent). Primary hepatocytes were seeded on collagen-coated 24-well Seahorse plates at 4 x 10^4^ cells/well and kept overnight in normal growth medium. Next day, cells were rinsed twice and kept in XF base medium (Seahorse Bioscience; Agilent) supplemented with 0.2% fatty acid-free BSA, 10 mM glucose and 2 mM pyruvate, pH 7.4, for 1-2 hrs before the run. Mitoparaquat (mitoPQ) was added in port A at the indicated concentrations followed by 1 μM FCCP in port B and 1 μM rotenone/antimycin A in port C. Hepa 1-6 cells were seeded at 3 x 10^4^ cells/well and kept in normal growth medium until adhering. Next, DMEM without glucose (Sigma, MO) supplemented with 20 mM galactose, 4 mM pyruvate, 2 mM glutamine and 10% CCS was added, and cells were kept overnight. The following day, cells were rinsed twice and kept in XF base medium supplemented with 0.3% fatty acid-free BSA with 20 mM galactose, 4 mM pyruvate, 2 mM glutamine, pH 7.4, for 1-2 hrs before the run. S1QEL 2.2 (ChemDiv, Inc) was added at the indicated concentrations in port A. For all experiments basal OCR was monitored for 15 min and mitoPQ and S1QEL 2.2 effects were plotted as % change of baseline in vehicle injected wells.

OCR of AML12 and Hepa 1-6 cells expressing Aox were measured using a Clark-type electrode fitted in a water-jacketed chamber (Strathkelvin Instruments, Scotland, UK). Cells were permeabilized with 0.02% digitonin in KHE buffer supplemented with 2 mM MgCl_2_, 10 mM KH_2_PO_4_, 0.3 % w/v fatty acid-free BSA, pH 7.1 at 37°C. Recording started in the presence of 5 mM succinate, 1 μM rotenone and 1 mM ADP. At the indicated time points potassium cyanide (KCN,100 µM) and n-propylgallate (50 μM) were injected into the chambers to inhibit complex IV and Aox, respectively.

### NADH redox microscopy

Primary hepatocyte NAD(P)H autofluorescence was detected by exciting at 340/26 nm and the signal emitted was collected using a DAPI cube (460/80 nm). The exposure time was 50 ms every 20 s for a total of ∼15 min. Hepatocytes were imaged in DMEM without glucose and at the indicated time points 5 mM pyruvate, 2 µM FCCP and 1 µM rotenone were added. Approximately 40 cells were analyzed per dish and the results are an average of at least 4 mice per condition. We report NADH fluorescence as NAD(P)H because NADH and NAD(P)H fluorescence properties are indistinguishable. However, the majority of the signal obtained in isolated mitochondria and intact cells is attributed to NADH autofluorescence because the NAD-pool is much larger. NADH signal is enhanced when bound to complex I and we use mitochondrial substrates/inhibitors to specifically modulate the NADH signal.

### Overexpression of alternative oxidase

Ciona intestinalis AOX (accession #: XP_018672179.1) gene sequence was codon-optimized for mammalian expression (IDT Codon Optimization Tool). The codon optimized AOX with a C-terminal HA tag synthesized by IDT was cloned into pLv-EF1a-IRES-puro (Addgene 85132), pAAV.TBG.PI.eGFP.WPRE.bGH (Addgene 105535), and pAdv-MCS-IRES-GFP (Vector Biolabs), respectively, for the production of viral particles. The adeno-associated virus (AAV) was produced by the Penn Vector Core. The adenovirus was produced by the Vector Biolabs.

AML12 and Hepa 1-6 cells were infected with lentiviral particles collected from the media of HEK293T cells transiently transfected with pLv-EF1a-AOX-HA-IRES-puro, psPAX2 and pMD2.G. Cells with AOX-HA stable expression were selected by treatment with puromycin (2 μg•mL^−1^) for 2 weeks.

### CoQ_10_ emulsion

A total of 100 mg of CoQ_10_ (Sigma-Aldrich) and corn oil (2:1, w/w) were dissolved in 1mL chloroform/methanol (2:1, v/v). Next, 150 µL of sodium oleate (2 mg•mL^−1^) and 500 µL phosphatidylcholine (Avanti Polar lipids) (25 mg•mL^−1^, in chloroform) were mixed together. The mixture was dried under argon for 30 minutes at room temperature and desiccated under a vacuum overnight at 4°C. The next day, the viscuous mixture was warmed at 60°C for 5 minutes and mixed with pre-warmed 0.9 mL 2% glycerol in PBS with 0.25 mM EDTA, pH 8.4. The solution was then sonicated, 1-second pause/on, for 10 mins at 50°C. The sonicated emulsion was centrifuged for 30 seconds to remove impurities. The CoQ_10_ emulsion was stored in a glass vial at room temperature. CoQ_10_ concentration was determined using the molar extinction coefficient ε_275_ = 14.6 mM^−1^cm^−1^ (CoQ_10_), and 10mg•kg^−1^ was injected every other day for 4 weeks.

### *In vivo* treatments

Male mice aged 9-12 weeks (23-34g) were acutely treated with mitoparaquat (MitoPQ, Cayman Biosciences) via intraperitoneal injection (i.p.) 1-2 hours before experimentation. MitoPQ was titrated from 1-6 nmol/mouse, which is equivalent to 0.04-0.2 mg•kg^−1^ body weight. Experiments were performed with 0.16 mg•kg^−1^ body weight (4 nmol/mouse) to avoid mitoPQ toxicity. Male mice aged 5 weeks were treated with the CoQ_10_ (10 mg•kg^−1^ body weight) or control emulsion via i.p. injection every other day for up to 9 weeks. Lactate/pyruvate tolerance tests from 2 cohorts were performed at 4, 7 and 9 weeks of age and the results were averaged.

### Glucose, lactate/pyruvate, insulin, and glycerol tolerance tests

For all tolerance tests animals were fasted for 16 hrs overnight, except for insulin tolerance tests where animals were daytime fasted for 6 hrs. For glucose tolerance tests, lean mice acutely treated with MitoPQ or DMSO vehicle received an i.p. injection of 1g•kg^−1^ glucose. Obese mice expressing aav.Aox or aav.GFP or chronically treated with CoQ_10_ emulsion or a control emulsion, received an i.p. injection of 0.5 g•kg^−1^ glucose. For the insulin tolerance tests, lean mice received an i.p injection of 0.7 U•kg^−1^ insulin (Humulin R U-100, Lilly, USA) while obese mice received 1.5 U•kg^−1^. Insulin was prepared in PBS containing 0.2% fatty acid-free BSA. For lactate/pyruvate tolerance tests lean mice acutely treated with mitoPQ received an i.p. injection of 1.5 mg•kg^−1^ of a solution containing 10:1 lactate to sodium lactate dissolved in PBS at 500 mg•mL^−1^. For glycerol tolerance tests lean mice acutely treated with mitoPQ received an i.p. injection of 1 g•kg^−1^ glycerol. All substrates were injected in a volume of ∼250μL diluted in PBS. Blood glucose and lactate levels were monitored before mitoPQ injection (time −60 min), just before substrate injection (time 0 min) and for up to 2 hrs following injection using a glucometer (Bayer Contour Next EZ) and a lactate meter (Nova Biomedical Lactate Plus) from a superficial nick at the tip of the tail.

### *In vivo* insulin signaling

Mice were fasted for 16 hrs and then received 4nmol mitoPQ or DMSO vehicle via i.p. injection. Two hours later mice were were anesthetized with 100 mg•kg^−1^ ketamine and 10 mg•kg^−1^ xylazine and then injected with 0.75 U•kg^−1^ of insulin (HumulinR, Lilly), prepared as described above, into the portal vein. Livers were collected 3 min later and immediately frozen in liquid N_2_. Epididymal adipose tissue and gastrocnemius were next collected and frozen until protein extraction. Obese mice expressing aav.Aox or aav.GFP received 3.5 U•kg^−1^ of insulin.

### Histology

Tissues were fixed in 10% zinc formalin overnight and then transferred to 70% ethanol for prolonged storage. Tissue processing, sectioning, and staining with hematoxylin and eosin and periodic acid-Schiff (PAS) for glycogen detection were performed by Histowiz. Adipocyte area was analyzed using QuPath image analysis software. Three randomly selected ROIs (2mm x 2mm) were analyzed per tissue slide. An adipose detector classifier was created, which applied a morphological closing function with a smoothing sigma filter setting of 0.5 and a threshold of 240. Individual adipocytes were then detected, and several morphometric parameters were exported (area, length, circularity, solidity, max diameter and min diameter). Positive PAS staining intensity was identified using the Halo image analysis software from Indica Labs and areas were quantified using the algorithm v2.1.3. The algorithm was set-up to measure two stains by first defining the settings to identify the magenta PAS stain and the blue heamatoxylin counterstain. The algorithm uses a color deconvolution step to separate the two stain colors, and this is then followed by setting thresholds for the PAS stain to detect weak, moderate and strong (Halo threshold settings 0.3876, 0.6199, 0.9518) staining. Data was expressed as percent of total area positive for PAS stain.

### MitoSOX oxidation

Oxidation of MitoSOX (Invitrogen) was used as indicator of superoxide levels in primary hepatocytes. MitoSOX oxidation was measured after 30 min incubation with 1 µM MitoSOX in DMEM without phenol red and supplemented with pyruvate and 0.2% fatty acid-free BSA, according to the manufacturer’s specifications.

### Glucose production

Primary hepatocytes were isolated and maintained in Williams E medium supplemented with 0.1 % CCS overnight. The next day cells were washed in warmed PBS and incubated in DMEM without phenol red and glucose, with 0.2 % fatty acid-free BSA in the presence of the gluconeogenic substrates 2 mM pyruvate, 2 mM glutamine and 20 mM lactate, or 20 mM glycerol. After 4h, medium was collected for measurement of glucose levels and the cells were washed and harvested for protein assay. Glucose was measured using 5 U•mL^−1^ glucose oxidase, 50 µM Amplex Ultra Red and 10 U•mL^−1^ HRP after 30 min incubation. Fluorescence was monitored using a microplate reader (SpectraMax Paradigm, Molecular Devices) at Λ_excitation_= 530 nm, Λ_emission_= 590 nm. Fluorescence was calibrated using glucose as standard.

### H_2_O_2_ release and insulin signaling from primary hepatocytes

Primary hepatocytes from obese mice were isolated as described above. Cells were seeded (2×10^4^) in a 96 well-plate and 4 hours after platinh they were washed and incubated overnight in Williams E media while exposed to adenoviral particles expressing Aox or GFP at a multiplicity of infection (MOI) of 100. For the H_2_O_2_ production assay, cells were washed in assay buffer [120mM NaCl, 3.5 mM KCl, 1.8 mM CaCl_2_, 0.4mM KH_2_PO_4_, 20 mM TES, 5mM NaHCO_3_, 1.2 mM Na_2_SO_4_, 1mM MgCl_2_] supplemented with 0.1% fatty acid-free BSA. The detection system consisted of 0.2 U•ml^−1^ HRP, 50 μM Amplex UltraRed, 100 μM PMSF, 24 U•ml^−1^ SOD and pyruvate and glutamine were used as substrates. S1QEL 2.2 (2.5-5 µM) was used to determine site I_Q_ contribution and H_2_O_2_ generated via RET was calculated by subtracting the rate of H_2_O_2_ generation with substrate alone from the rate in the presence of S1QEL. Fluorescence was monitored using a microplate reader (SpectraMax Paradigm, Molecular Devices) and was calibrated using H_2_O_2_ as standard. For insulin signaling, primary hepatocytes (the day after isolation) were incubated in serum-free Williams E media for 4hrs. 3-18nM of insulin was then added to the media and after 3 min cells were washed twice with 1X PBS and immediately frozen in liquid N_2_.

### Protein extraction and immunoblotting

Liver, epidydimal fat and skeletal muscle were homogenized with a TissueLyser II in NP-40 buffer [50 mM Tris-HCl (pH 7.4), 2 mM EGTA, 5 mM EDTA, 30 mM NaF, 10 mM Na_3_VO_4_, 10 mM Na_4_P_2_O_7_, 40 mM glycerophosphate, 1 % NP-40, and 1% protease inhibitor cocktail] supplemented with 10 nM okadaic acid. Homogenates were incubated for 10 min on ice and then centrifuged at 14000 g for 10 min. Supernatant was collected and protein concentrations were determined by BCA method using albumin as a standard. Samples were diluted in 6x Laemmli loading buffer and heated at 95°C for 5 min. Protein was separated by 4-12% NU-PAGE gradient gels using 1x MOPS buffer (Invitrogen), or 4-20% Criterion TGX-stain free gels (BioRad) and using 7.5 % SDS-PAGE using 1x Tris/glycine-SDS buffer (BioRad). Blots were incubated with primary antibody overnight, washed 3 times and then incubated with HRP-conjugated secondary antibody (Cell Signaling Technologies) for 1h at RT. Membranes were visualized by chemiluminescence (Roche Diagnostics) using a BioRad ChemiDoc MP imaging system.

### Endogenous protein staining and confocal imaging

Primary hepatocytes were seeded on 35 mm round glass bottom imaging dishes in Williams Medium with 5% CCS and maintained overnight at 37°C, 5% CO_2_. The following morning, cells were washed and incubated in DMEM supplemented with 2mM pyruvate and glutamine and 50 nM MitoTracker^TM^ Deep Red (Invitrogen) for 45 min and then fixed with 4% paraformaldehyde for 10 min at RT. The cells were then washed 3x in PBS and permeabilized for 20 min with 0.2% Triton-X100 in PBS at RT. Anti-HA primary antibody (#3724, Cell Signaling Technologies) was used to visualize aav.Aox-HA and was diluted 1:200 in PBS, and the cells were incubated in this solution overnight at 4°C. The next day, cells were washed 3x with PBS, including one long wash for more than 10 min. Cells were then incubated with 1:1000 diluted secondary antibody in PBS for 1h at RT in the dark. The cells were washed 3x, including one long wash. Cells were imaged with a Yokogawa CSU-X1 spinning disk confocal system (Andor Technology, South Windsor, CT) with a Nikon Ti-E inverted microscope (Nikon Instruments, Melville, NY), using a 60x or 100x Plan Apo objective lens with a Zyla cMOS camera, and NIS elements software was used for acquisition parameters, shutters, filter positions and focus control. Image analysis was performed using Fiji software. At least 30 cells from each genotype were analyzed.

### Primary and secondary antibodies

Primary antibodies used were, <u>anti-beta tubulin</u> (ab21058), <u>anti-NDUFS1</u> (ab169540), <u>anti-VDAC</u> (ab14734), <u>oxphos cocktail</u> (MS604; ab110413) from Abcam. <u>Anti-ND6</u> (A32848) was from Boster Biological Technology. <u>Anti-phospho-insulin receptor (IR)</u> (pTyr972) (I1783-1VL) was from Sigma. <u>Anti-NDUFV2</u> (SC-271620), <u>anti-insulin receptor</u> <u>β</u> (SC-711), <u>anti-NDUFA6</u> (SC-86755) were from Santa Cruz Biotech. <u>Anti-phopho-AKT</u> (pThr308) (cs-9275), <u>anti-pan AKT</u> (cs-4691), <u>anti-phospho-GSK-3α/β</u> (cs-9331), <u>anti-HA</u> (cs-2367), <u>anti-calreticulin</u> (cs-12238), <u>anti-pJNK</u> (cs-4668), <u>anti-SAPK/JNK</u> (cs-9252) were from Cell Signaling Technologies. <u>Anti-GSK-3α/β</u> (44610) was from Thermo Fisher Scientific. Secondary antibodies used were, anti-rabbit IgG-HRP (7074), anti-mouse IgG-HRP (7076), and anti-rabbit IgG Alexa Fluor 488 (4412) from Cell Signaling Technologies.

### Gene expression

RNA was isolated from liver tissues from 9–10-week-old wildtype and ob/ob mice. Tissues were homogenized in TRIzol (Invitrogen) using a TissueLyser II (Qiagen) at 30 Hz for 3 min at 4°C. RNA was extracted using Nucleospin RNA kits (#740955.250, Macherey-Nagel). Complementary DNA was synthetized using iScript RT Supermix kit (BioRad). Quantitative PCR (qPCR) was run in triplicates on a ViiA7 RT-PCR System (Applied Biosystems) using SYBR green and custom primers or primer sets based on sequences from the Harvard Primer Bank. The cycle thresholds (Cts) of target genes were normalized to hypoxanthine phosphoribosyltransferase (HPRT1) Cts to calculate expression levels using the 2^−ΔΔCt^ method. For the genes in the mevalonate and CoQ biosynthetic pathway, we used the average of HPRT CTs between 4 independent experiments to normalize the CTs of the target genes.

### Primer list

<u>Coq2</u> (L- GACCCAGGTTGTTTTCCAGA; R- TGGAAGGTCGAAATGTCTCC), <u>coq3</u> (L- TCGTGGCTTCTGAAGTTGTG; R- CCCACGTATGAGTGCCTTTT), <u>coq5</u> (L- CCACGGTCTGTGACATCAAC; R- GCCTGGTCAATGTGTGTGAC), <u>coq7</u> (L- TTTGGACCATAGCTGCATTG; R- ATGCGGTTTGCTCCATATTC), <u>coq8a</u> (L- GAAGTCTGGGCTGCAGTAGG; R- GAAGCCTGCCTTTTTGTCTG), <u>coq9</u> (L- CCTGTCAAAATCCCCTGAGA; R- GCTCAGCACAACTGTCCAAA), <u>pdss1</u> (L- CCGCGACTTTCAGACTTGA; R- TTGGGGAGGCAGACATTAAA), <u>pdss2</u> (L TGGTGCATCGTGGGATAGTA; R- TACTCCATGCACCAAGTCCA), <u>HMGCS primer set 1</u> (L- CTCTGTCTATGGTTCCCTGGCT; R- TCCAATCCTCTTCCCTGCC), <u>HMGCS primer set 2</u> (L- TGATCCCCTTTGGTGGCTGA; R- AGGGCAACGATTCCCACATC), <u>HMGCR1</u> (L- CCGGCAACAACAAGATCTGTG; R- ATGTACAGGATGGCGATGCA), <u>HMGCR2</u> (L- ATCCTGACGATAACGCGGTG; R- AAGAGGCCAGCAATACCCAG), <u>MVK</u> (L- CTCTGCTTGCCTTTCTCTAC; R- TCGGGAGTGTCCTGAAATA), <u>PMVK</u> (L- CTGTTTAGCGGGAAGAGAAA; R- GGCAGAGCTACATCTTCATAG), <u>MVD</u> primer set 1 (L- AAGCAGACGGGCAGTACAGT; R- CCTGGAGGTGTCATTGAGGT), <u>MVD primer</u> set 2 (L- CTGCACCAGGACCAGCTAAA; R- CTGAGGCTGAGGGGTAGAGT), <u>IPP isomerase</u> (L- GACGTCAGGCTTGTGCTAGA; R- CTAGAACACAGAGATTCCGGCT), <u>FPP synthase</u> (L- TCCAGGTCCAGGACGACTAC; R- CGCCTCATACAGTGCTTTCA); <u>GGPP synthases</u> (L- GACAAGCTACAGATTATCATTGAAGTG; R- ATCCGGGTGATCAAGGGTTA), and <u>HPRT1</u> (L- CCAGCGTCGTGATTAGCG; R- CCAGCAGGTCAGCAAAGAAC).

### Body composition, Comprehensive Lab Animal Monitoring System (CLAMS)

Body composition of 7 week old ob/ob mice 14 days after aav.Aox or aav.GFP injection was measured using magnetic resonance imaging (EchoMRI™, TX, USA). For CLAMS, 7 week old ob/ob mice expressing aav.AOX or aav.GFP for 12 days were housed individually and acclimatized for 1 day. Oxygen consumption, carbon dioxide release, energy expenditure (EE) and activity were measured using a Columbus Instruments Oxymax-CLAMS system as previously described, according to guidelines for measuring energy metabolism in mice (*57*).

### LC/MS conditions

For small polar metabolite separation and data acquisition in extracted tissues, a Vanquish Horizon (Thermo Fisher Scientific, Waltham, MA) ultra-high liquid chromatography (LC) system coupled to a Thermo Q Exactive HF Orbitrap mass spectrometer (MS) was used. For separation, a Waters (Milford, MA) XBridge BEH Amide (2.5 μm, 2.1×150 mm) column fitted with a VanGuard (2.5 μm, 2.1×5 mm) guard column was used. The mobile phases were as follows: Phase A: 95% water/5% acetonitrile and Phase B: 20% water/80% acetonitrile with 10 mM ammonium acetate and 10 mM ammonium hydroxide in both phases.

The flow rate was held constant at 0.3 ml/min and the following gradient conditions were used: 0 min, 100% B; 3 min, 100% B; 3.2 min, 90% B; 6.2 min, 90% B; 6.5 min, 80% B; 10.5 min, 80% B; 10.7 min, 70% B; 13.5 min, 70% B; 13.7 min, 45% B; 16 min, 45% B, 16.5 min, 100% B; and 22 min, 100% B. The samples were held at 4°C, the injection volume was 5 μl and the column was held at 25°C.

The separated metabolites were analyzed in both positive and negative ionization modes in the same run (switching mode). The mass spectra were acquired using a resolution of 120,000 in the 70 – 1,000 m/z range. The ElectroSpray Ionization source parameters for both modes were as follows: capillary temperature 300°C, spray voltage 3.5kV, sheath gas 40 (arbitrary units), auxiliary gas 10 (arbitrary units), probe heater temperature 30°C and S-Lens RF level 45 v.

### Human study subjects

Patients with overweight or obesity and non-alcoholic fatty liver disease (NAFLD) from the Liver Unit of Hospital Clínic in Barcelona were studied. The study had been approved by the Hospital Clinic IRB (HCB/2019/0458). All patients gave written informed consent, and two specimens of liver biopsy were collected, one for histopathological diagnosis and one for microarray analysis (n=14) or CoQ determination (n=13). Approximately 2 mg liver biopsies for CoQ measurements were immediately frozen in liquid N_2_ upon collection. Clinical, demographic and laboratory data were collected in all subjects at the time of liver biopsy and NAFLD was diagnosed according to clinical guidelines (*61*, *62*). Steatosis and fibrosis grade were defined according to the NASH CRN criteria (*63*). Healthy subjects within the microarray cohort were liver donors from the Liver Unit Transplant Program in the Hospital Clinic from which liver biopsies were obtained at the time of living-donor liver transplantation. Characteristics of patients at the time of inclusion are included in Supplementary Tables 1 and 2.

### Patient sample mRNA isolation and gene expression analysis

mRNA from the transcriptome-cohort was isolated from fresh human liver tissues using TRIzol, according to the manufacturer’s instructions (Invitrogen, Carlsbad, CA, USA). RNA samples were hybridized to GeneChips and Affymetrix Human Genome U219 arrays. All microarray data used in the present study was deposited and it is accessible on the public repository of NCBI Gene Expression Omnibus (GEO) (GSE139602 and GSE224645). In this manuscript we only analyzed the subset of genes involved in the CoQ biosynthesis.

### Statistics and Reproducibility

Statistical significance was assessed using GraphPad Prism Version 7. All data is mean ± SEM of n independent values. Student’s t-test was used to calculate statistical significance between two groups. For multiple comparison tests one-way ANOVA using Tukey’s or Dunnett’s correction was applied. Where genotype/treatment versus time were analyzed, 2-way ANOVA followed by Šidák’s post-hoc test was used and when appropriate Dunnett’s correction was applied. P < 0.05 was considered statistically significant.

**Table S3.**
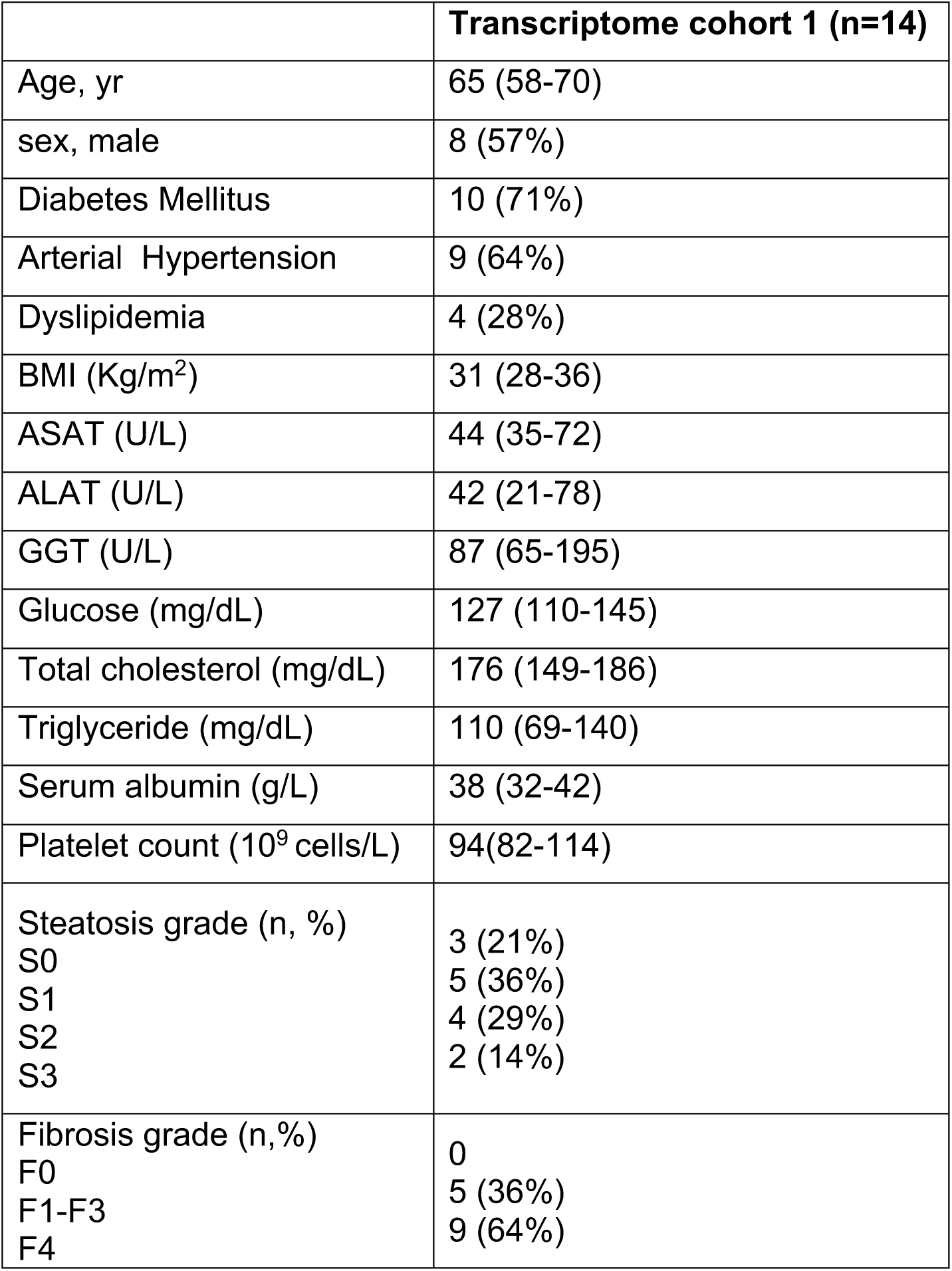
Characteristics of subjects undergoing liver biopsies for gene expression analysis.

**Table S4.** Characteristics of subjects undergoing liver biopsies for CoQ redox state analysis.

|  | **Cohort 2 CoQ redox state (n=13)** |
| --- | --- |
| Age, yr | 64 (55-65) |
| Sex, male | 12 (92%) |
| Diabetes Mellitus | 2 (15%) |
| Arterial hypertension | 4 (31%) |
| Dyslipidemia | 5 (38%) |
| BMI (Kg/m<sup>2</sup>) | 32 (28-36) |
| ASAT (U/L) | 40 (22-52) |
| ALAT (U/L) | 46 (23-65) |
| GGT (U/L) | 44 (31-77) |
| Glucose (mg/dL) | 104 (94-120) |
| Total cholesterol (mg/dL) | 164 (115-198) |
| Triglyceride (mg/dL) | 120 (89-211) |
| Serum albumin (g/L) | 46 (41-47) |
| Platelet count (10<sup>9</sup> cells/L) | 224 (136-239) |
| Steatosis grade (n, %) |  |
| S0 | 2 (15%) |
| S1 | 7 (53%) |
| S2 | 3 (23%) |
| S3 | 1 (7.7%) |
| Fibrosis grade (n,%) |  |
| F0 | 4 (31%) |
| F1-F3 | 5 (38%) |
| F4 | 4 (31%) |

---

**Fig. S1.**
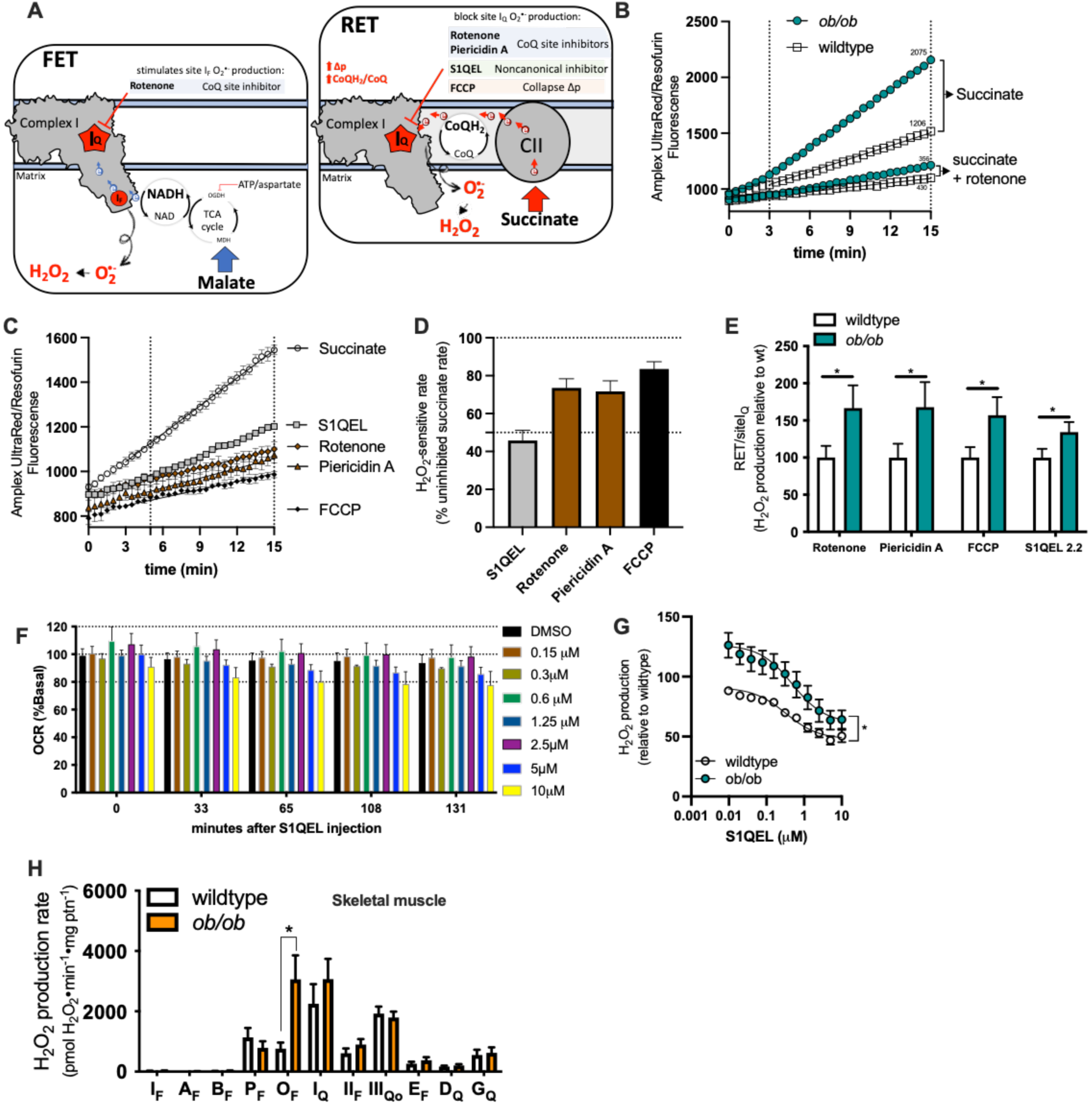
ROS generation by RET from site I_Q_ is increased in the liver but not the skeletal muscle of obese mice. (**A**) Schematic to illustrate the substrates and inhibitors used to assess the two modes of mROS generation from complex I: forward electron transport from site I_F_ (FET, left) and reverse electron transport (RET, right) from site I_Q_. (**B**) Representative Amplex UltraRed traces showing that rotenone blocks mROS during succinate oxidation. These rates difference define RET, which is higher in mitochondria isolated from ob/ob livers. (**C**) Representative traces of Amplex UltraRed oxidation during succinate oxidation in the presence or absence of 5 µM S1QEL 2.2, 2 µM rotenone, 2 µM piericidin A, and 1µM FCCP in isolated mitochondria. (**D**) Quantification of (C). (**E**) Relative mROS production by RET, from ob/ob liver mitochondria, as defined by the compounds in (D). **(F**) Effect of S1QEL 2.2 (0.15-10 µM) on Hepa 1-6 oxygen consumption rates (OCR). (**G**) mROS production during RET from liver-isolated mitochondria from wildtype and ob/ob mice treated with different concentrations of S1QEL 2.2 (0.01-10 µM). (**H**) Maximum capacity of superoxide/H_2_O_2_ production from skeletal muscle-isolated mitochondria from lean wildtype and ob/ob mice (WT and ob/ob 8-10 weeks of age). Values are means ± SEM. *, p<0.05; **, p<0.01; ***, p<0.0001 by Student’s *t* test.

**Fig. S2.**
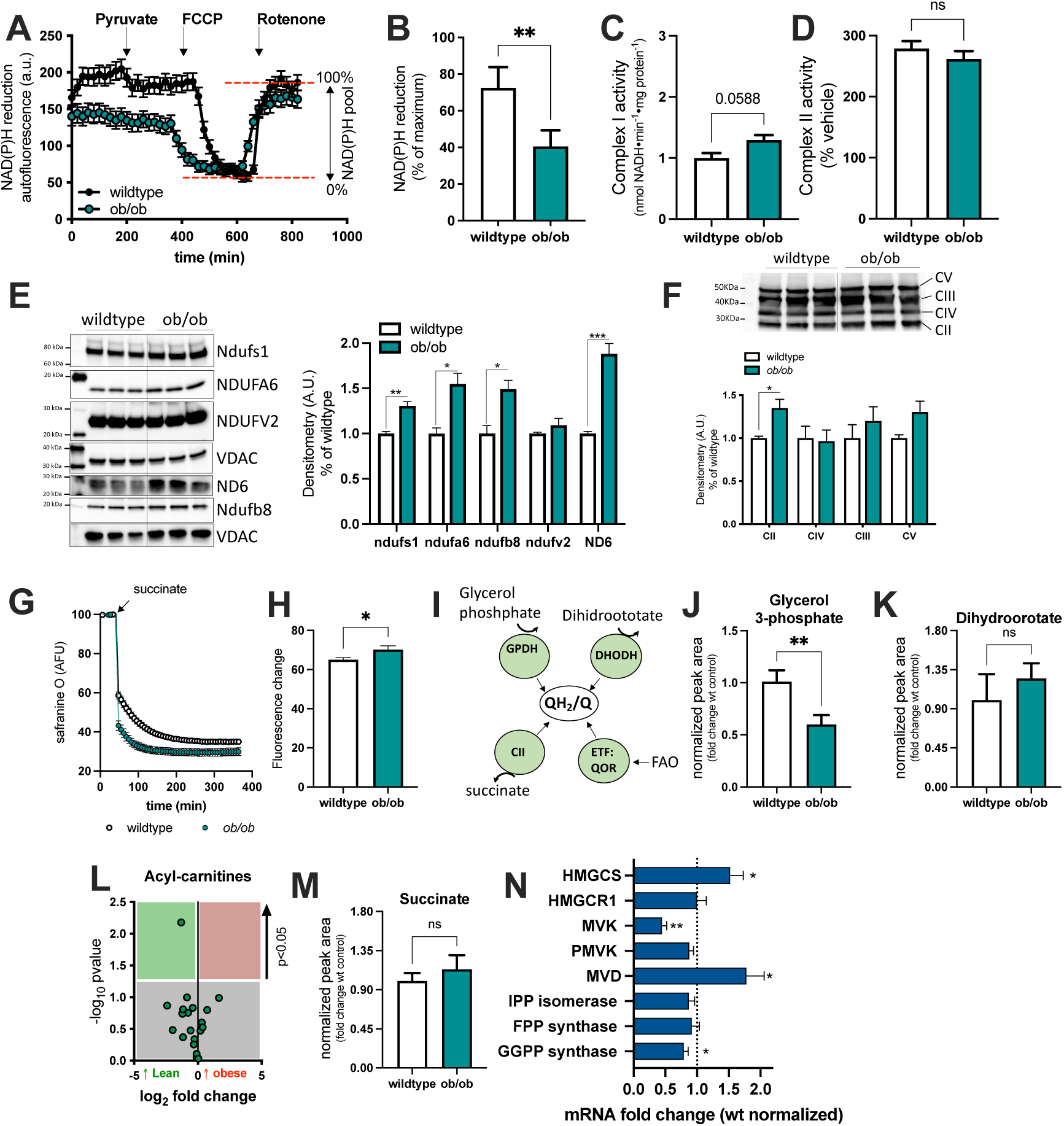
The thermodynamic forces driving mROS via RET. (**A**) Kinetics of NAD(P)H autofluorescence during the oxidation of 5 mM pyruvate in primary hepatocytes isolated from wildtype (wt) and ob/ob mice. The NAD(P)H pool was oxidized after addition of 1µM FCCP (0% NAD(P)H) and the 100% value was established by the addition of 2µM rotenone. (**B**) Percent reduction of NAD(P)H during pyruvate oxidation. (**C**) Complex I and (**D**) complex II activity in wt and ob/ob isolated mitochondria. (**E**) Immunoblot (left) and quantification analysis (right) of complex I subunits in the livers of wt and ob/ob mice. (**F**) Immunoblot (top) and quantification analysis (bottom) of complex II-V of the electron transport chain (ETC) in the livers of wt and ob/ob mice. (**G**) Mitochondrial membrane potential from wt and ob/ob livers was measured with safranine O during the oxidation of 5 mM succinate plus 4µM rotenone in the presence of oligomycin. (**H**) Quantification of wt and ob/ob mitochondrial membrane potential. (**I**) Illustration of the enzymes that shuttle electrons into the CoQ pool that generate mROS. Quantification of the levels of (**J**) glycerol phosphate, (**K**) dihydroorotate, (**L**) acyl-carnitines and (**M**) succinate in the livers of wt and ob/ob mice. (**N**) Relative expression levels of the genes in the mevalonate pathway in the livers ob/ob mice relative to wt. Values are means ± SEM. *, p<0.05; **, p<0.01; ***, p<0.001; ****, p<0.0001 by Student’s *t* test.

**Fig. S3.**
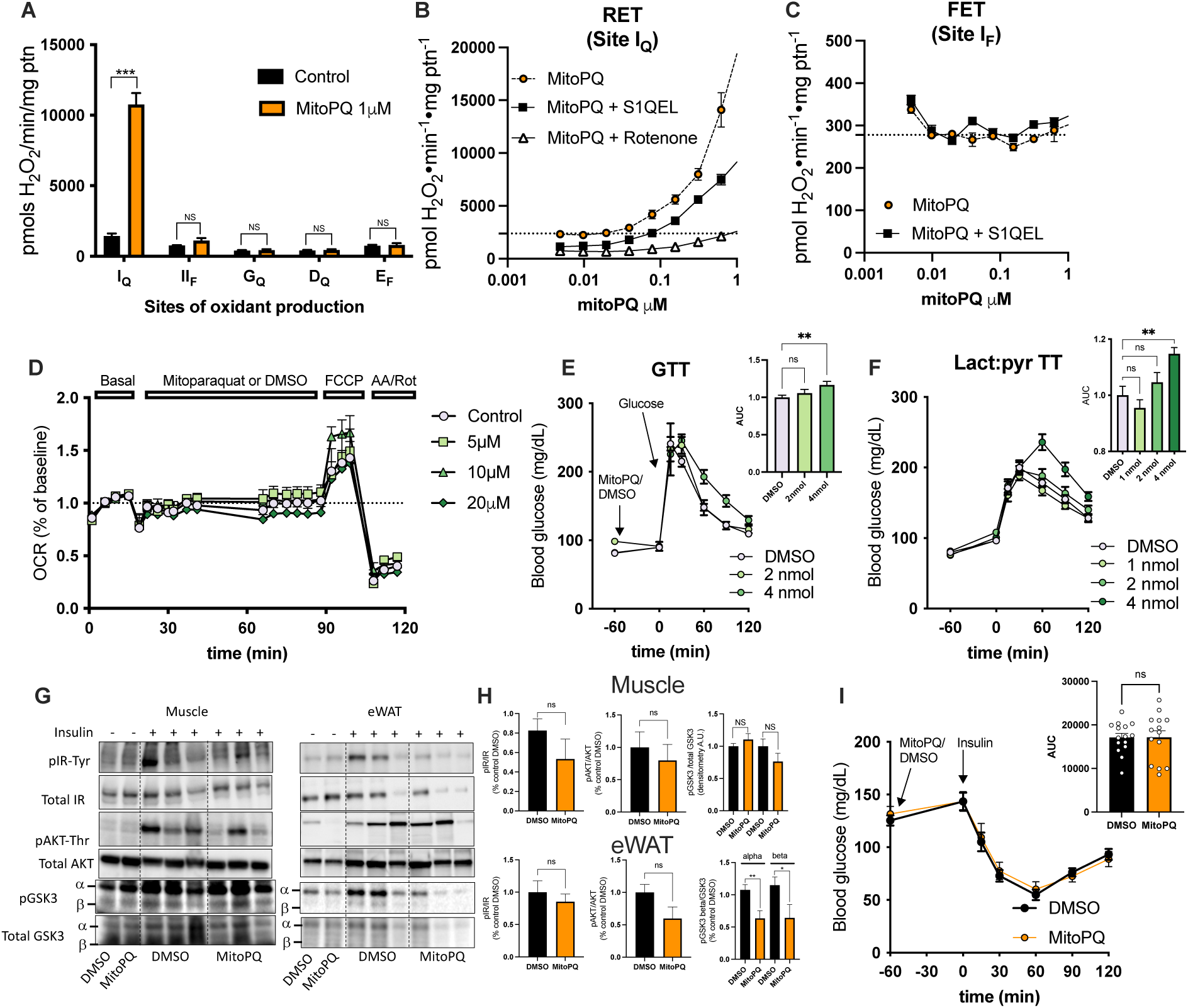
Mitoparaquat promotes mROS via RET and impairs glucose homeostasis in the liver. (**A**) Effect of 1 µM mitoPQ on superoxide/H_2_O_2_ production from the sites linked to the Q-pool (sites I_Q_, II_F_, G_Q_, D_Q_ and E_F_) (n=3 independent mitochondrial isolations). (**B**) Effect of mitoPQ ± 10 µM S1QEL 2.2 or 2 µM rotenone on the rate of superoxide/H_2_O_2_ production by RET or (**C**) FET. (**D**) Effect of mitoPQ on the oxygen consumption rate (OCR) of wildtype (wt) primary hepatocytes. MitoPQ was injected in port A and OCR was monitored for the next hour. 1 µM FCCP and 2 µM rotenone and antimycin A were injected in ports B and C, respectively (n=2 preparations per group). (**E**) Blood glucose levels during i.p. glucose tolerance test (0.5 g•kg^−1^) in mice treated with 2-6 nmol mitoPQ. (**F**) Blood glucose levels during i.p. lactate: pyruvate tolerance test (1.5 : 0.15 g•kg^−1^) in mice treated with 1-4 nmol mitoPQ. (**G**) Immunoblot of *in vivo* insulin signaling in the gastrocnemius muscle (left) and epididymal fat (right) of wt mice 1.5 h after 4 nmol mitoPQ or DMSO treatment (n=8 mice per group). (**H**) Immunoblot analysis. (**I**) Blood glucose levels during insulin tolerance test (0.7 U insulin•kg^−1^) in 6 hour-fasted mice treated with 4 nmol mitoPQ or DMSO. *Inset*: area under the curve. Values are means ± SEM. **, p<0.01; ***, p<0.001; ****, p<0.0001 by Student’s *t* test.

**Fig. S4.**
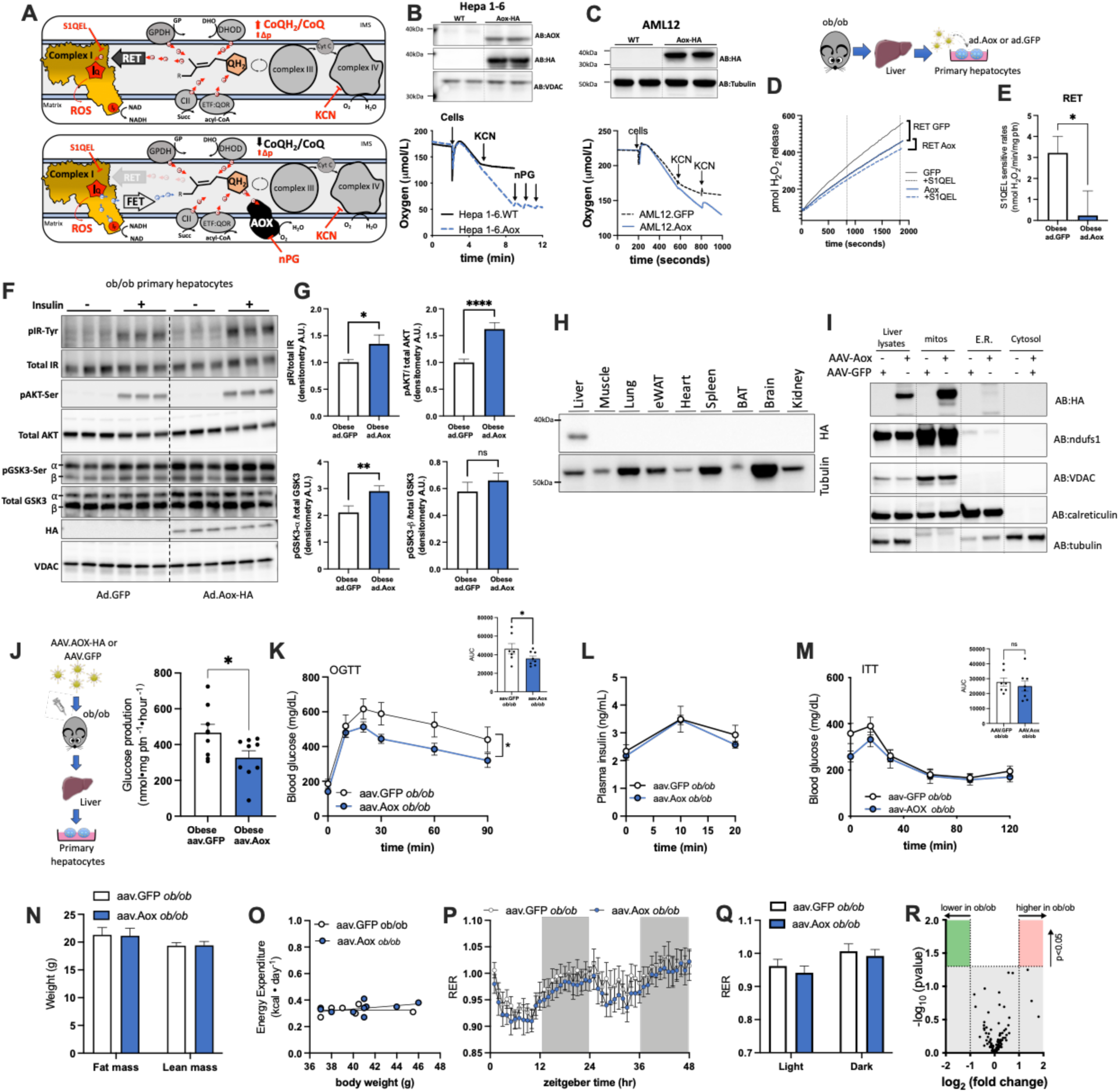
Ectopic expression of Aox in hepatocytes decreases mROS generation via RET and improves systemic glucose homeostasis. (**A**) Illustration showing mROS production in ob/ob mice ± Aox expression. Aox decreases mROS by RET. (**B**) Representative traces of cyanide insensitive oxygen consumption in Hepa 1-6 and (**C**) AML12 expressing Aox. N-propyl gallate inhibits Aox activity. *Top,* immunoblot confirming Aox expression. (**D**) Amplex UltraRed traces to show that Aox-expressing primary hepatocytes generates less mROS by RET. Ad., adenovirus. (**E**) Rate of H_2_O_2_ release by RET measured as the S1QEL sensitive rate. (**F**) Immunoblot of 18 nM insulin action in isolated hepatocytes from 9-10 week old obese mice incubated with ad.Aox-HA or ad.GFP for 24h. (**G**) Immunoblot analysis. (**H**) Immunoblot from tissue homogenates of Aox expressing mice. (**I**) Immunoblot from different cellular fractions from the liver of GFP or Aox mice. ndufs1 and VDAC, mitochondria; calreticulin, endoplasmic reticulum and tubulin, cytosol. (**J**) Glucose production from primary hepatocytes isolated from obese mice expressing Aox or GFP using 20 mM lactate, 2 mM pyruvate, and 2 mM glutamine as substrates. (**K**) Blood glucose levels during oral glucose tolerance test (OGTT) (0.75 g•kg^−1^) in ob/ob mice. *Inset,* area under the curve. (**L**) Plasma insulin levels during OGTT. (**M**) Blood glucose levels during insulin tolerance test (3.5 U of insulin•kg^−1^). *Inset*, area under the curve. (N-R) Metabolic profile of obese 10 days after Aox expression. (**N**) body composition of 7 week old ob/ob mice following aav administration. (**O**) Energy expenditure (EE) as a function of bodyweight. (**P**) Respiratory exchange ratio measured during metabolic cage housing. (**Q**) Analysis of RER during light and dark cycles. (**R**) Metabolite levels in the livers of ob/ob mice expressing Aox vs. GFP. Values are means ± SEM. *, p<0.05; **, p<0.001; ****, p<0.0001 by Student’s *t*-test or multiple comparisons analysis by 2-way ANOVA followed by Sidak’s post-hoc analysis.

**Fig. S5.**
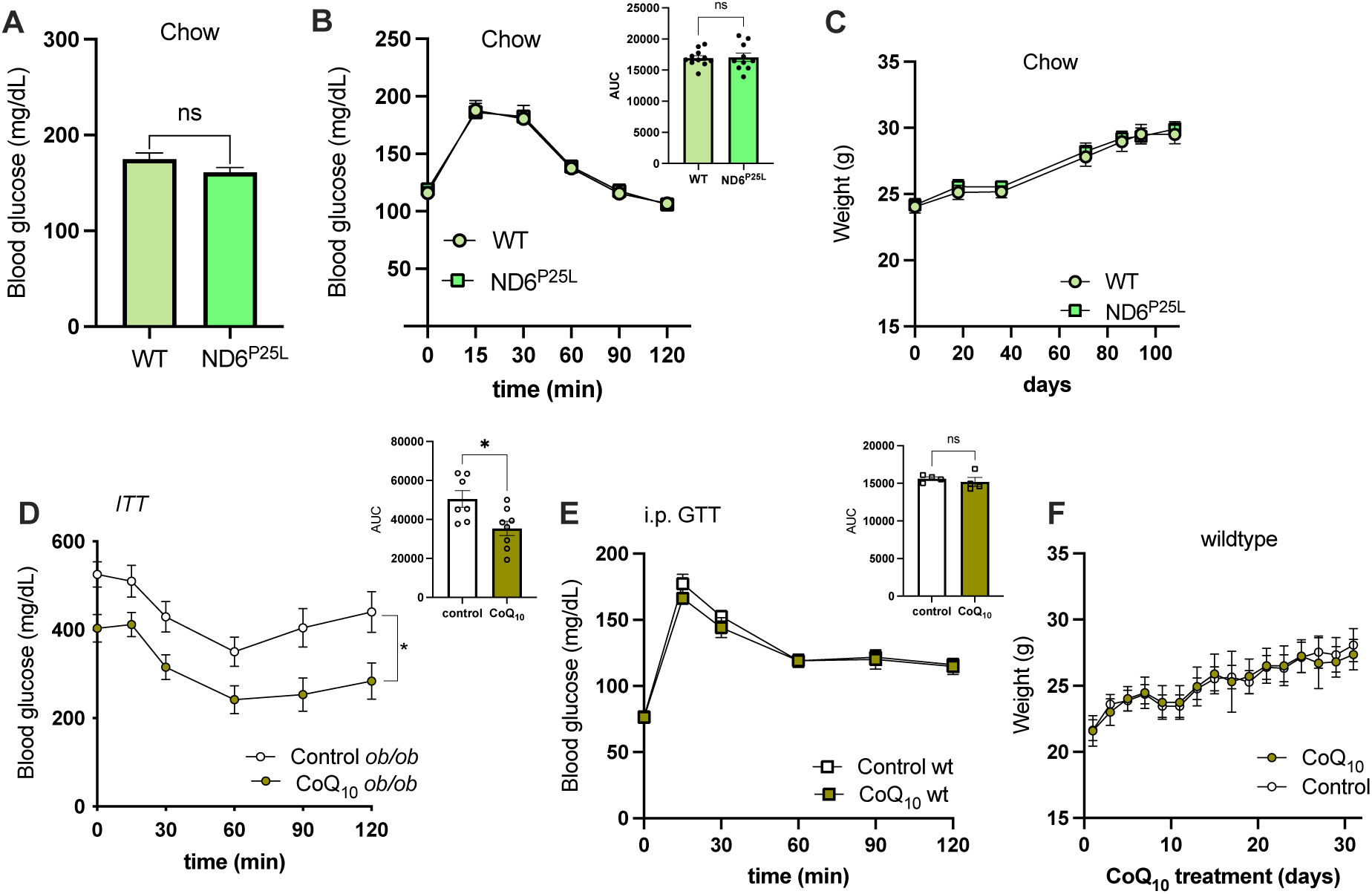
Suppressing RET in lean mice does not change body weight or glucose tolerance. (**A**) Six hour fasting blood glucose levels in wildtype and ND6^P25L^ mice fed chow diet for 15 weeks. (**B**) Blood glucose levels during glucose tolerance test (0.5 g•kg^−1^) in *ND6*^P25L^ and wildtype mice on chow diet for 10 weeks. *Inset,* area under the curve. (**C**) Weight gain of wildtype and ND6^P25L^ mice after 15 weeks on chow. (**D**) Blood glucose levels during insulin tolerance test (1.5 U of insulin•kg^−1^) in 6 hour fasted ob/ob mice treated with 10mg•kg^−1^ CoQ_10_ or vehicle every other day for 23 days. *Inset,* area under the curve. (**E**) Blood glucose levels during glucose tolerance test (0.5 mg glucose /g body weight, i.p) in wildtype mice treated with 10mg•kg^−1^ CoQ_10_ or vehicle every other day for 23 days. *Inset,* area under the curve. (**F**) Bodyweights of wildtype mice treated with CoQ_10_ or vehicle. Values are means ± SEM. *, p<0.05; by Student’s *t* test or multiple comparisons analysis by 2-way ANOVA followed by Sidak’s post-hoc analysis. Table S1. Substrates and inhibitors used to measure site-specific superoxide production in isolated mitochondria.

